# Müller glia subtypes define neuro-glial associations and spatial morphogen axes in the zebrafish retina

**DOI:** 10.64898/2026.03.22.713558

**Authors:** Samuel S Storey, Carrie L Hehr, Shaelene Standing, Sarah McFarlane

## Abstract

Müller glia are instrumental macroglia of the vertebrate retina, once thought to be a homogeneous population. Now Müller glia are generally accepted as transcriptionally heterogeneous, and new evidence suggests functional diversity may exist in the way these cells respond to retinal injury. It remains unclear, however, whether this functional heterogeneity is limited to a transient phenotype that stems from injury or a fundamental feature of the healthy retina. Here, we investigate Müller glia heterogeneity in the uninjured zebrafish retina across development and adulthood using a comprehensive single-cell transcriptomic atlas of the 5 days post-fertilization (dpf) eye, validated *in vivo* and integrated with 9 dpf and adult datasets. We reveal three constitutive subpopulations of Müller glia that persist from early larval stages into adulthood: 1) a proliferative and immature population in both the peripheral and central retina; 2) a novel cohort of neuron-associated Müller glia that express coherent transcriptional programs specific to distinct neuronal subtypes, including retinal ganglion, amacrine and horizontal cells; and 3) spatially distinct Müller glia subsets that appear to define a dorso-ventral axis of retinoic acid metabolism, bisected by a novel *cyp26c1*-expressing equatorial domain. Finally, cross-species analysis reveals that while neuron-associated programs are evolutionarily conserved in mammals, the spatial patterning of morphogens in adult retinae may be specific to the teleost lineage. Collectively, these findings provide robust evidence for intrinsic functional heterogeneity in the uninjured vertebrate retina, reframing Müller glia from a general support population to a specialized cellular network that actively maintains retinal geography and function.

**Main points:**

- Zebrafish Müller glia are heterogeneous in early larval retina.
- Zebrafish Müller glia subtypes define a spatial axis of retinoic acid metabolism.
- Neuron-associated glial programs identified in Zebrafish Müller glia are evolutionarily conserved in mammals.

## Introduction

The vertebrate retina is a specialized extension of the central nervous system that sits at the back of the eye where it converts photic signals into visual images through complex cellular interactions. While the neural retina is organized into three distinct nuclear laminae including the ganglion cell layer (GCL), the inner nuclear layer (INL) and the outer nuclear layer (ONL), the computational power of this tissue relies on intricate vertical columns of functional neural circuitry. Photoreceptors in the ONL convert light into electrical signals and synapse with bipolar cells in the INL that in turn relay the signal to retinal ganglion cells in the GCL. Horizontal and amacrine interneurons provide secondary regulation of synaptic transmission at the outer and inner synaptic (plexiform) layers, respectively (reviewed in Hoon et al., 2014; Masland, 2001). To explain the sophistication of this processing, research has cataloged an immense taxonomy of neuronal subtypes. Kolb et al. (2001) described almost 60 unique cell subtypes in the vertebrate retina, and subsequent work has characterized over 150 neuronal subtypes in the chick retina alone (Yamagata et al., 2021). However, while we increasingly appreciate this neuronal diversity, similar efforts have often overlooked the non-neuronal elements of retinal physiology.

Müller glia are the predominant macroglia of the vertebrate retina and embody structural, neurotrophic, homeostatic, synaptogenic and neurogenic roles to serve as the functional core of retinal circuitry columns (reviewed in Reichenbach & Bringmann, 2013). Anatomically, Müller glia extend vertical processes that span the full thickness of the retina and horizontal processes that interact with cells both neighboring and afar (MacDonald et al., 2015). Functionally, they recycle, synthesize, and coordinate the release of neurotransmitters and neurotrophic factors (Bringmann et al., 2006), and maintain metabolic, ionic, and water homeostasis (Tsacopoulos & Magistretti, 1996). The hallmark function of Müller glia is their reaction to injury or disease. In mammals, Müller glia respond to severe retinal trauma or disease with the formation of a glial scar, while in select teleosts like the zebrafish, Müller glia respond to cell death by complete functional regeneration (Bringmann et al., 2009; Karl & Reh, 2010; Nagashima et al., 2013; Nelson et al., 2012; Powell et al., 2016; Ramachandran et al., 2010). Further, zebrafish Müller glia feature a lifelong homeostatic neurogenic phenotype, being the sole source of new rod photoreceptors in the postnatal eye (Bernardos et al., 2007; Kustermann et al., 2010), with all other cell types made by a peripheral neurogenic niche called the ciliary marginal zone (CMZ) (Karl & Reh, 2010; reviewed in Hoon et al., 2014; reviewed in Lahne et al., 2020). Despite these multifaceted roles, the functional heterogeneity of Müller glia in the vertebrate retina remains poorly characterized.

There is strong evidence to suggest that retinal Müller glia, much like their neuronal counterparts, represent a transcriptionally heterogeneous population. As the transcriptome strongly predicts cellular function, this transcriptional heterogeneity implies functional heterogeneity. Early single-cell RNA sequencing (scRNAseq) experiments pointed to transcriptional heterogeneity in Müller glia of the chick (Yamagata et al., 2021) and human (Menon et al., 2019) retina. More recently scRNAseq analyses in the zebrafish demonstrate transcriptional heterogeneity, and suggest that functional heterogeneity exists within the regenerative context as some, but not all, larval and adult Müller glia become reactive under injury conditions (Celotto et al., 2023; Krylov et al., 2023; H. Xu et al., 2025). This implies a fundamental functional distinction between neurogenic and non-neurogenic Müller glia. The question remains whether functional heterogeneity is limited to a transient phenotype that stems from injury or a fundamental feature of the healthy retina.

Here, we investigate Müller glia heterogeneity in the uninjured context across three key stages of retinal development. We use single-cell transcriptomics and *in vivo* validation in the 5 days post-fertilization (dpf) larval zebrafish eye to demonstrate that Müller glia are a diverse cell type with robust heterogeneity beginning as early as 5 dpf. We focus here on three functional archetypes: 1) a novel cohort of neuron-associated glia, 2) spatially restricted subtypes defined by morphogen metabolism, and 3) peripheral and central niches of proliferative and undifferentiated cells. We integrate publicly available datasets from 9 dpf (Krylov et al., 2023) and adult (6-12 months post fertilization; Celotto et al., 2023) retina to demonstrate that these functional categories are constitutively maintained, and that Müller glia diversity persists throughout the lifespan. Further, we perform a cross-species analysis with public datasets from the E18 chick (Yamagata et al., 2021), post-embryonic mouse (Li et al., 2024), fetal human (Kriukov et al., 2025; Lu et al., 2020; Sridhar et al., 2020) and adult human (Li et al., 2026) to reveal that key features of Müller glia heterogeneity, particularly the neuron-associated programs and morphogen metabolism, are evolutionarily conserved in mammals.

Collectively, these findings indicate that Müller glia can be resolved into distinct populations. Our data suggests this compartmentalization contributes to the broad establishment of spatially segregated retinal microenvironments and precise, cell-specific regulation by glia-neuron interactions. These insights reframe Müller glia from a uniform population into specialized, integral components of the neural retina in vertebrates and lifelong retinal neurogenesis in the teleost.

## Methods

### Zebrafish Husbandry and Transgenic Lines

Zebrafish (*Danio rerio*) embryos and larvae were maintained in E3 according to standard procedures on a 14h light/10h dark cycle at 28.5°C. All experimental protocols were approved by the University of Calgary Animal Care Committee. Wild-type embryos were of the Tüpfel longfin (TL) background. The *Tg(gfap:egfp)* line was used to visualize Müller glia morphology and to validate cell identity in immunohistochemical assays. Larvae were staged by days post-fertilization (dpf) (Xu et al., 2020).

### Single-cell Dissociation and Library Preparation

Whole retinas were processed to capture the full diversity of retinal cell types without the potential bias or loss of fragile cells associated with cell sorting. Approximately 40,000 5 dpf wild type zebrafish retinal cells were harvested for scRNAseq over two independent collections. The protocol was modified from Xu et al. (2020). Eyes from 5 dpf larvae were dissected in E3 with tricaine and collected on ice. Eyes were then dissociated into a single-cell suspension using activated papain (Worthington) for 15 minutes at 37 °C with gentle mechanical trituration every 2-3 minutes. The enzymatic reaction was stopped using wash buffer (Xu et al., 2020) and cells were filtered through a 30µm strainer to remove clumps. Cells were spun down and resuspended to a final volume of 50µL in phosphate buffered saline (PBS) containing 0.2% Bovine Serum Albumin (BSA, Sigma Aldrich) and 1µL RNaseOUT (Invitrogen). Viability was assessed for resuspended cells using trypan blue and a hemocytometer. Approximately 10,000 cells were loaded onto the Chromium Single Cell Chip (10x Genomics). Single-cell droplet encapsulation and library preparation were performed using the 10x Genomics Chromium Next GEM Single Cell 3’ Reagent Kit (v3.1) according to the manufacturer’s instructions. Libraries were sequenced on an Illumina NovaSeq 6000 to a depth of 11,027 counts per cell.

### Bioinformatic Analysis

Raw sequencing data were aligned to the zebrafish genome (GRCz11_v4.3.2) using Cell Ranger (v3.1.0, 10x Genomics). Downstream analysis was performed using the Seurat R package (v4.3.0, Hao et al., 2021). For the initial retinal cell atlas, cells were filtered to remove low-quality droplets based on unique molecular identifier (UMI) counts (1,683 median UMI per cell; 741 median genes per cell; removed cells with greater than 2500 or less than 200 genes) and mitochondrial transcript percentage (>5%). To annotate technical doublets, doublet scores were estimated per sample with Scrublet (Wolock et al., 2019) using an expected doublet rate of 0.06 and a fixed score threshold of 0.20 to yield a predicted doublet rate of 2.77%. Data were normalized using the LogNormalize method and batch correction was achieved using CCA integration. Dimensionality reduction was performed using Principal Component Analysis (PCA) on highly variable genes. The top 30 principal components were used to construct a K-nearest neighbor (KNN) graph, and clusters were identified using the FindClusters function (resolution = 0.8). Non-linear dimensional reduction for visualization was performed using Uniform Manifold Approximation and Projection (UMAP). Cell types were annotated based on the expression of canonical marker genes derived from Hoang et al. (2020) and other established literature. The 5 dpf Müller glia subset was bioinformatically isolated based on the robust expression of *glula*, *rlbp1a*, and *apoeb*. This subset was re-clustered using the top 50 principal components (PCs). Differentially expressed genes (DEGs) for each cluster were identified using the FindAllMarkers function (Wilcoxon Rank Sum test, adjusted p-value < 0.05). To quantify neuronal gene signatures, the AddModuleScore function in Seurat was used with gene lists derived from the top 50 DEGs of annotated neuronal clusters in the full atlas. RNA velocity analysis was performed using scVelo (Bergen et al., 2020) in Python after conversion of the SeuratObject (R) to AnnData (Virshup et al., 2024) using the scCustomize package (Marsh et al., 2025). Spliced and unspliced reads were counted from .loom files using the velocyto package (La Manno et al., 2018) and velocity vectors were projected onto the pre-computed UMAP embeddings. For lifespan analysis, 5 dpf data were integrated with publicly available datasets from 9 dpf (Krylov et al., 2023) and adult (Celotto et al., 2023) retinas using CCA integration.

### Cross-species bioinformatics analysis

The pre-processed chick Müller glia dataset was downloaded from the Broad Institute Single Cell Portal and converted to an AnnData object in Python; chick data was used as-is, as the raw counts were not included in the csv data. Mouse, fetal human and adult human datasets were accessed from the CELLXGENE platform (CZI Single-Cell Biology et al., 2023). Bioinformatic isolation of Müller glia from the mouse, fetal human and adult human atlases was performed using standard scanpy subset methodology from ‘cell_type’ annotations inherent to the CELLXGENE schema v.7.0.0. QC validation for mitochondrial count and ribosomal associated protein count was evaluated on raw counts to ensure fidelity with the chick and our zebrafish data, and 2000 highly variable genes were calculated for each dataset using standard methods in scanpy (Wolf et al., 2018). Intra-dataset integration was performed with scVI (Gayoso et al., 2022) to correct for batch effects with ‘donor_id’ as the batch key. Leiden clustering was performed with a resolution of 0.6 for the comparison. We mapped zebrafish genes to orthologs (Supplemental Table 3) obtained from Ensembl BioMart for the chicken, mouse and human.

### Tissue Preparation and Sectioning

For histological analysis, larvae and adult eye cups were fixed in 4% paraformaldehyde (PFA) in PBS overnight at 4 °C. Following fixation, tissues were cryoprotected in 35% sucrose/PBS overnight, embedded in optimal cutting temperature (OCT; Sakura Tissue-Tek), and frozen. For ages 5 dpf to 9 dpf, four to seven fish were included per slide for fluorescent in situ hybridization. For 28 dpf samples, three to four fish were used per slide. Cryosections were cut at 12 µm thickness using a MICROM cryostat (HM 500 OM) and transferred to a slide.

### Riboprobe design

Digoxigenin (DIG)-labeled antisense riboprobes were synthesized from primers designed using the NCBI Primer Blast program for *her4.1, pcna, gad2, gad1b, onecut2, stc2a, rdh10a, aldh1a3, cyp26c1, bambia,* and *aldh1a2*. Briefly, reverse primers were ordered with T7 (TAATACGACTCACTATAGG) added to the 5’ end. *her4.1* F: AGTTCATCAAGCAGCAGCCC, R: GTAAGAGTACAGGCAATTCTTCTCC. *pcna* F: AGTGACGGGTTCGACTCCTA, R: GGACGTGTCCCATGTCTGCAAT. *gad2* F: CTGCATGAGTTCAGTGAC, R: GCAGTGGAGGTTAGCCAAT. *onecut2* F: GCCCGAGTTTCAGAGGATGT, R: GCAAAACTGTGAACTGCTGGGT. *stc2a* F: TGATGTGCACGAGAGTCACC, R: CTTCGAGTCTTCCTGCTCGG. *rdh10a* F: AAAGTGCGTAGGGAGGTTGG, R: ACACGATTGATTCAAACGGC. *aldh1a3* F: GAGTCGAAGGACACTGGCAA, R: ATGTACGTGGCTCCATTGGG. *cyp26c1* F: AGCACACGAATCATCCTGGG, R: GCCATCAGTGATCAAGCCCT. *bambia* F: GCCACGGGGTATCAAGACAA, R: CATTCGCAACGCCAGCATAA. *aldh1a2* F: CTGAACGGGGGAAGCTACTG, R: AGGTCCGTGTTCAGTGGTTG.

### *in situ* Hybridization (ISH)

RNA whole mount (WM) ISH and fluorescent in situ hybridization (FISH) were conducted using standard protocols (Kalifa et al., 2025; Thisse & Thisse, 2008). Briefly, Digoxigenin (DIG) labelled RNA probes were generated by PCR-amplification; the 5’ end of the reverse primers contained the DNA sequence for T7 RNA polymerase (T7: 5′-GAAATTAATACGACTCACTATAGG-3′). For whole mount ISH, fixed, methanol-dehydrated embryos were rehydrated with PBS-Tween 20 (PBST) washes and permeabilized with proteinase K followed by post-fixation with 4% PFA. Permeabilized embryos were hybridized overnight at 65 °C in hybridization buffer containing the DIG-labelled RNA probe. High-temperature washes at 65 °C removed unhybridized probe. For immunodetection, tissues were blocked in a solution of PBST supplemented with 2% sheep serum and 2mg/mL bovine serum albumin (BSA) and incubated overnight at 4 °C with alkaline phosphatase-conjugated anti-digoxigenin antibody diluted 1:1000.

The chromogenic reaction was developed at room temperature using nitro blue tetrazolium (NBT) and 5-bromo-4-chloro-3-indolyl phosphate (BCIP) substrate, and embryos were post-fixed in 4% PFA prior to imaging. For FISH on frozen sections, a similar protocol was followed pursuant to the following modifications: 1) samples were stained for 30 minutes in the dark with fluorescein tyramide at 1:50 in 1x Amplification Buffer (Akoya BioSciences) containing 2% dextran sulfate, 2) RNA probes were added directly to the cryostat sections on glass slides and coverslipped prior to incubation. For every marker, riboprobe expression was confirmed in at least four separate embryos.

### Immunohistochemistry

Sections were washed in PBS containing 0.8% Triton X-100 and blocked in wash buffer with the addition of 1% BSA for one hour at room temperature. Primary antibodies were applied overnight at 4°C. The following primary antibodies were used: mouse anti-Glutamine Synthetase (1:400; Millipore), rabbit anti-GFP (1:500; Molecular Probes), mouse anti-PCNA (1:300; Sigma-Aldrich), rabbit anti-Calbindin (1:100; Swant), and mouse anti-HuC/D (1:100; Molecular Probes). Following washing, species-specific secondary antibodies conjugated to Alexa Fluor 488, 568, or 647 (1:500; Invitrogen) were applied for one hour at room temperature. Nuclei were counterstained with Hoechst (Invitrogen).

### Microscopy and Image Analysis

Fluorescent images were acquired using a Zeiss LSM 900 laser scanning confocal microscope using a 20x objective and ZenBlue software. Whole mount images were acquired by brightfield microscopy using a dissecting compound microscope (Zeiss Axiocam-HRc stereoscope). Image processing, including brightness and contrast adjustments and orthogonal projections, was performed using FIJI/ImageJ (Schindelin et al., 2012). Figure assembly was performed using PowerPoint.

## Results

### Single cell RNA sequencing reveals Müller glia transcriptional heterogeneity in the larval retina

To define the transcriptional landscape of Müller glia during functional maturation we generated a single-cell RNA sequencing (scRNAseq) atlas of the whole retina at 5 days post-fertilization (dpf), when the retina is functionally developed and the zebrafish exhibits visually evoked behaviours (Easter & Nicola, 1996; Fadool & Dowling, 2008). We used common sets of transcribed genes (Hoang et al., 2020) to resolve and annotate all cell types of the neural retina, other supporting ocular tissues (retinal pigment epithelium [RPE], lens, vascular tissue, cornea), and the neurogenic ciliary marginal zone (CMZ) (Karl & Reh, 2010; reviewed in Hoon et al., 2014; reviewed in Lahne et al., 2020). Figure 1A demonstrates our resulting cell atlas of the 5 dpf retina with a Uniform Manifold Approximation and Projection (UMAP) plot.

**Figure 1.**
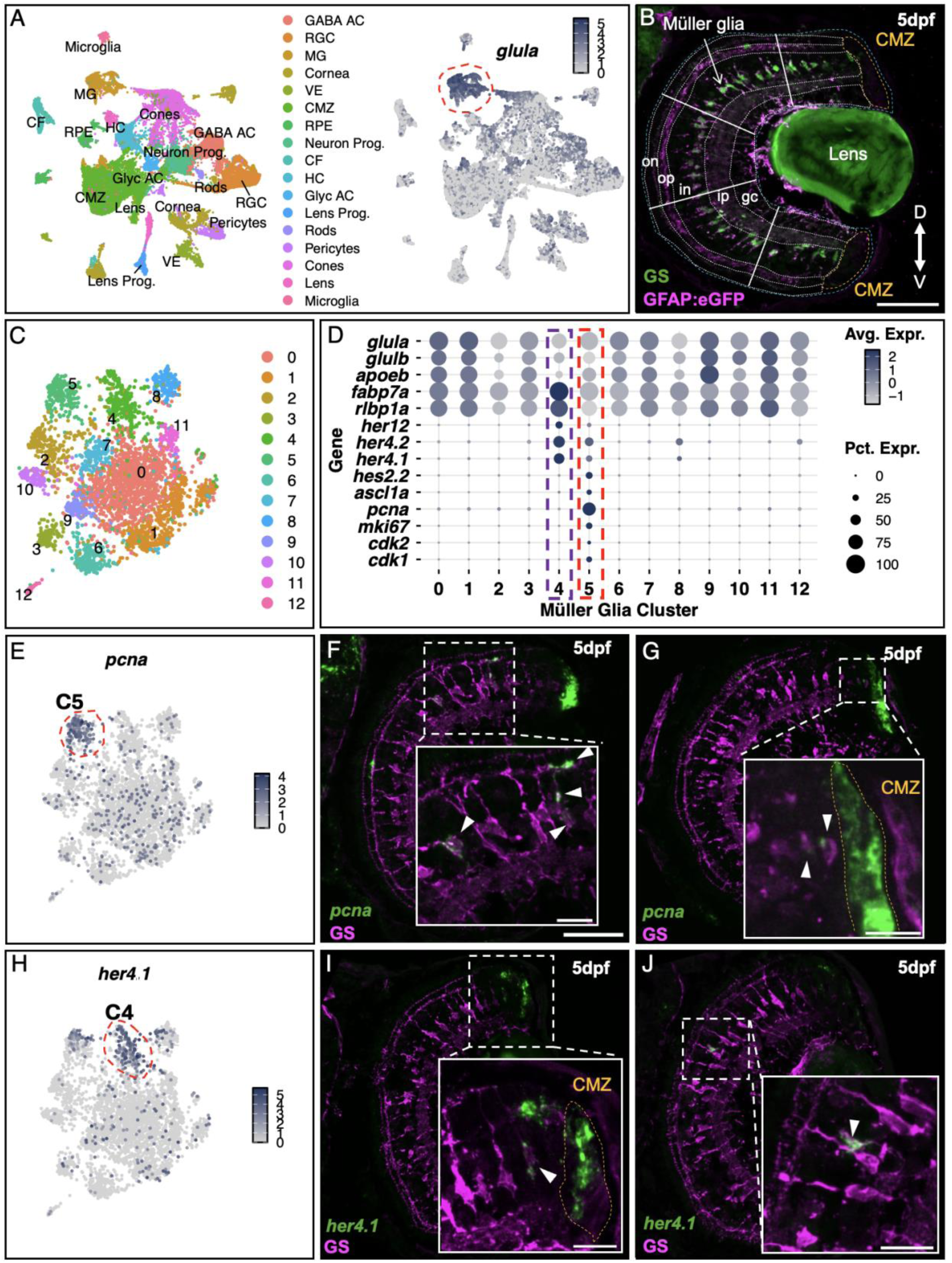
Müller glia are heterogeneous in the uninjured, wild type 5 dpf zebrafish retina. A) *Left:* Uniform manifold approximation plot (UMAP) of scRNAseq data from the whole zebrafish eye with annotated cell populations at 5 days post-fertilization (dpf). Abbreviations: GABA AC = GABAergic amacrine cell, RGC = retinal ganglion cell, MG = Müller glia, VE = vascular epithelium, CMZ = ciliary marginal zone, RPE = retinal pigmented epithelium, CF = corneal fibroblasts, HC = horizontal cells, Glyc AC = glycinergic amacrine cells. *Right:* UMAP detailing expression of *glutamine synthetase* mRNA (*glula;* GS) within the Müller glia population. B) Fluorescence immunohistochemistry (IHC) against Glutamine Synthetase (Glula; GS) in a transverse section of the neural retina of a 5 dpf *Tg(gfap:egfp)* zebrafish. Blue dashed lines delineate eye border. White dashed lines delineate laminar anatomy: on = outer nuclear layer, op = outer plexiform layer, in = inner nuclear layer, ip = inner plexiform layer, gc = retinal ganglion cell layer. The ciliary marginal zone (CMZ) is outlined with gold dashes. Solid white lines divide the retinal sections into five segments for spatial contextualization: the peripheral retina is defined as both the dorsal and ventral outer segments and includes the CMZ; the central retina spans the three inner segments; the equatorial retina constitutes the centre-most segment. Axes: D = dorsal, V = ventral. C) UMAP of 13 transcriptionally distinct clusters (C0-C12) identified by unbiased clustering of the 5 dpf Müller glia subset. D) Expression dot plot for core gene expression in the Müller glia subset. Dot size represents the percentage of cells within the cluster expressing the specific gene (Pct. Expr.), and the intensity of dot color represents the scaled, mean log-normalized expression level (Avg. Expr.). Dashed boxes highlight the differential enrichment of genes in clusters C4 (purple) and C5 (red). E, H) UMAP plot with expression of *pcna* (E; C5 encircled in red dashes) and *her4.1* (H; C4 encircled in red dashes) transcripts within the 5 dpf Müller glia subset. F, G, I, J) Fluorescent *in situ* hybridization (FISH; green) of 5 dpf transverse retinal sections alongside anti-GS immunostaining (magenta) to visualize transcript co-expression within Müller glia and their processes. White dashed boxes are enlarged with solid borders to show region of interest. GS and transcript double-positive Müller glia are shown by white arrowheads in enlarged section. Scale bars: F is 50 µm; F, G, I, J inset scale bars are 15 µm.

Within our unbiased clustering, the robust and specific co-expression of canonical markers of glial function identified Müller glia: glutamine synthetase (*glula/b*; GS; UMAP expression encircled with red dashed line Figure 1A) which clears synaptic glutamate; retinaldehyde binding protein (*rlbp1a*), a key mediator of the visual cycle; and apolipoprotein Eb (*apoeb*) involved in lipid metabolism and transport (Hoang et al., 2020; Lyu et al., 2023; Raymond et al., 2006; Saari et al., 2001; Schlegel et al., 2021; H. Xu et al., 2025). UMAP expression plots for key Müller glia markers in the whole eye atlas are demonstrated in Supplementary Figure S1.

*In vivo,* double fluorescence immunohistochemistry (IHC) against GS and GFP on a *Tg(gfap:egfp)* background confirmed that mature Müller glia cell bodies and processes in the 5 dpf retina expressed membrane-bound GS and glial fibrillary acidic protein (GFAP) (Bernardos et al., 2007; Raymond et al., 2006) (Figure 1B). To clearly contextualize these spatial patterns, we divided the transverse retina cross-section into five segments; the peripheral retina is defined as the dorsal and ventral outer segments, and includes the ciliary marginal zone (CMZ) stem cell niche; the central retina spans the three inner segments; and the equatorial retina constitutes the centre-most segment (Figure 1B; spatial segments divided by orange dashed lines; CMZ encircled in gold dashes). Newly born Müller glia emerge from and abut the CMZ, and immature Müller glia transition to maturity within the peripheral segments; both populations weakly express GFAP and GS. Müller glia of the central retina expressed higher levels of GS. While high levels of *gfap*-driven eGFP accumulated and perdured in the central Müller glia, endogenous *gfap* mRNA exhibited low baseline expression (Bernardos et al., 2005; Celotto et al., 2023) and sparse capture in our uninjured scRNAseq dataset. Thus, we excluded *gfap* as a marker for bioinformatic isolation of Müller glia.

Müller glia constituted approximately 10% of the total captured cells (3,563 of 35,826 quality-control filtered cells) within our scRNA-seq dataset, an overrepresentation relative to the expected anatomical proportion of 4-5% (Lenkowski & Raymond, 2014). This overrepresentation likely reflects a recovery bias inherent to the dissociation protocol, which preferentially retains or recovers robust glial cells relative to more fragile neuronal cell types during tissue dissociation (Pollak et al., 2013).

We performed unsupervised clustering of the isolated Müller glia population with n=50 principal components and revealed thirteen transcriptionally distinct clusters (C0 – C12), as shown by UMAP in Figure 1C. We used the FindAllMarkers function in the Seurat R package (Hao et al., 2021) to identify differentially expressed genes (DEGs) within each Müller glia cluster (Supplementary Table 1) and suggest their potential biological function. Müller glia cluster C0 exhibited a relative paucity of differentially expressed markers and lack of subspecialized transcriptional programs but maintained high expression of canonical Müller glia genes (*glula/b, apoe, fabp7a, rlbp1a*) (Figure 1D). The remaining clusters displayed a remarkable heterogeneity, revealing a diversity of transcriptional programs layered upon a core Müller glia identity.

We first characterized Cluster 5 (C5), which we propose represents Müller glia in an active proliferative state. C5 was enriched for genes critical for cell cycle progression and proliferation, including *proliferating cell nuclear antigen* (*pcna*)*, marker of proliferation Ki-67* (*mki67*) (*Ki-67* in mammals), *cyclin-dependent kinase 1* (*cdk1*), *cyclin d1* (*ccnd1*), *achaete-scute family bHLH transcription factor 1a* (*ascl1a*) and *high-mobility group box 2b* (*hmgb2b*) (Bringmann et al., 2006, 2009; Celotto et al., 2023). These data are shown by dot plot in Figure 1D (red dashed box). The Seurat-generated dot plot combines percent expression (Pct. Expr.) to represent the proportion of cells within a specific cluster that express the gene, indicated by at least one unique molecular identifier (UMI), by dot size, and the average expression (Avg. Expr.) represents the mean log-normalized expression value for that gene across all cells in the cluster by intensity of colour, where darker values indicate greater expression. A UMAP for *pcna* enrichment in C5 is shown in Figure 1E (expressed in 11.3% Müller glia [403/3563 cells] by scRNAseq).

These data appear to map Müller glia across a developmental continuum spanning from immature (*her4.1/2*+, *fabp7a*+, *rlbp1a*++) to functional maturity (*glula+, cahz+, apoeb+*), and include actively cycling (*pcna+, cdk1+*) cells. To validate the C5 population *in vivo*, we performed fluorescent *in situ* hybridization (FISH) for *pcna* alongside GS immunostaining on 5 dpf retinal sections. We detected *pcna* expression in the occasional Müller glia cell of the mature central retina (Figure 1F, inset; >1 *pcna+* Müller glia observed in 106/106 central sections, n=6 larvae) consistent with the canonical neurogenic role of central retina Müller glia to produce rod photoreceptors (Bernardos et al., 2005; Campbell et al., 2022; Lenkowski & Raymond, 2014).

Additionally, we caught the occasional *pcna*+/GS+ cell in the peripheral retina next to the CMZ stem cell niche (Figure 1G; observed in 10/106 central retinal sections, n=6 larvae). We propose this latter group are either newly born Müller glia or Müller glia progenitors, which have just begun to express GS protein and not yet downregulated *pcna* mRNA. We performed immunohistochemistry with a PCNA antibody and confirmed the sporadic presence of Müller glia in the central eye that strongly express the protein (Supplemental Figure S2). Thus, the C5 population likely comprises proliferative Müller glia including both the neurogenic, rod progenitor-producing Müller glia of the central retina, and possibly progenitors of the peripheral retina that produce Müller glia for the new peripheral neural circuits made through the lifespan (Karl & Reh, 2010; Wan et al., 2016).

Cluster C4 lacked the proliferative markers that defined C5, but expressed similarly reduced levels of canonical markers of mature glia (*glula/b, apoeb*). Unlike the C5 population, C4 was most strongly enriched for *fatty-acid binding protein* (*fabp7a*), *rlbp1a*, and bHLH hairy-related Notch signaling targets (*her4.1, her4.2, her12, her15.1*) (Figure 1D, purple dashed box). A UMAP of *her4.1* localized to C4 is shown in Figure 1H (expressed in 9.6% Müller glia [343/3563]). FISH with a *her4.1* riboprobe showed infrequent colocalization with immature Müller glia that displayed GS immunoreactivity, specifically located adjacent to the CMZ (Figure 1I; observed in 16/41 central retinal sections, n=6 larvae; CMZ outlined in orange), and isolated Müller glia of the central retina (Figure 1J; observed in 5/41 central retinal sections, n=6 larvae). Established literature identifies *her4.1* as a Notch-responsive marker of reactive and progenitor-like Müller glia states (Mitra et al., 2018; Takke et al., 1999). Consequently, we hypothesize that C4 represents an immature, post-mitotic population, which could include either newborn Müller glia that arise from the CMZ (Raymond et al., 2006), or de-differentiated, neurogenic Müller glia of the central retina (Campbell et al., 2022; Celotto et al., 2023), or both.

### Several Müller glia subpopulations are defined by coherent neuronal gene programs

Unexpectedly, our DEG analysis revealed selective enrichment for a diversity of canonical neuronal lineage markers (Figure 2A) in Müller glia clusters C2, C3, C6, C7, and C12 as shown by expression dot plot in Figure 2B. Müller glia clusters C3, C6 and C12 were strongly enriched for the neuron-specific RNA binding protein marker *elavl3* (Figure 2B, top row). C3 cells were defined by an enrichment of retinal ganglion cell (RGC) fate determination markers, including the developmental transcription factor *isl2b,* neuritin family member *nrn1a* that is implicated in RGC synaptic plasticity and axonal outgrowth, and the POU-domain transcription factors (*pou4f1/2/3,* also known as *brn3a/b/c*) (Figure 2B, orange dashed box) (Badea et al., 2009; Gan et al., 1996; Isenmann, 2003; Sharma et al., 2015). C6 showed significant enrichment of gamma amino-butyric acid (GABA)-ergic amacrine cell markers including the GABA-synthesis enzymes, *glutamate decarboxylase 1b and 2* (*gad1b, gad2;* Figure 2B, red dashed box), and the GABA transporter critical for neurotransmitter reuptake, *slc6a1a* (Yan et al., 2020). C12 was the smallest Müller glia subpopulation, and strongly expressed markers associated with both GABA-ergic and glycinergic amacrine cell transcriptional programs, including the acetylcholine synthesis enzyme, *choline acetyl transferase* (*chata*; Figure 2B; green dashed box) (Chen et al., 2007). Notably, none of the neuron-associated clusters showed expression for the neuronal-differentiation factors *neurod1* or *neurod4* (Boutin et al., 2010; Hardwick & Philpott, 2015; Lenkowski & Raymond, 2014), or the pro-neuronal marker *olig2* (McFarland et al., 2008) (Figure 2B; bottom two rows).

**Figure 2.**
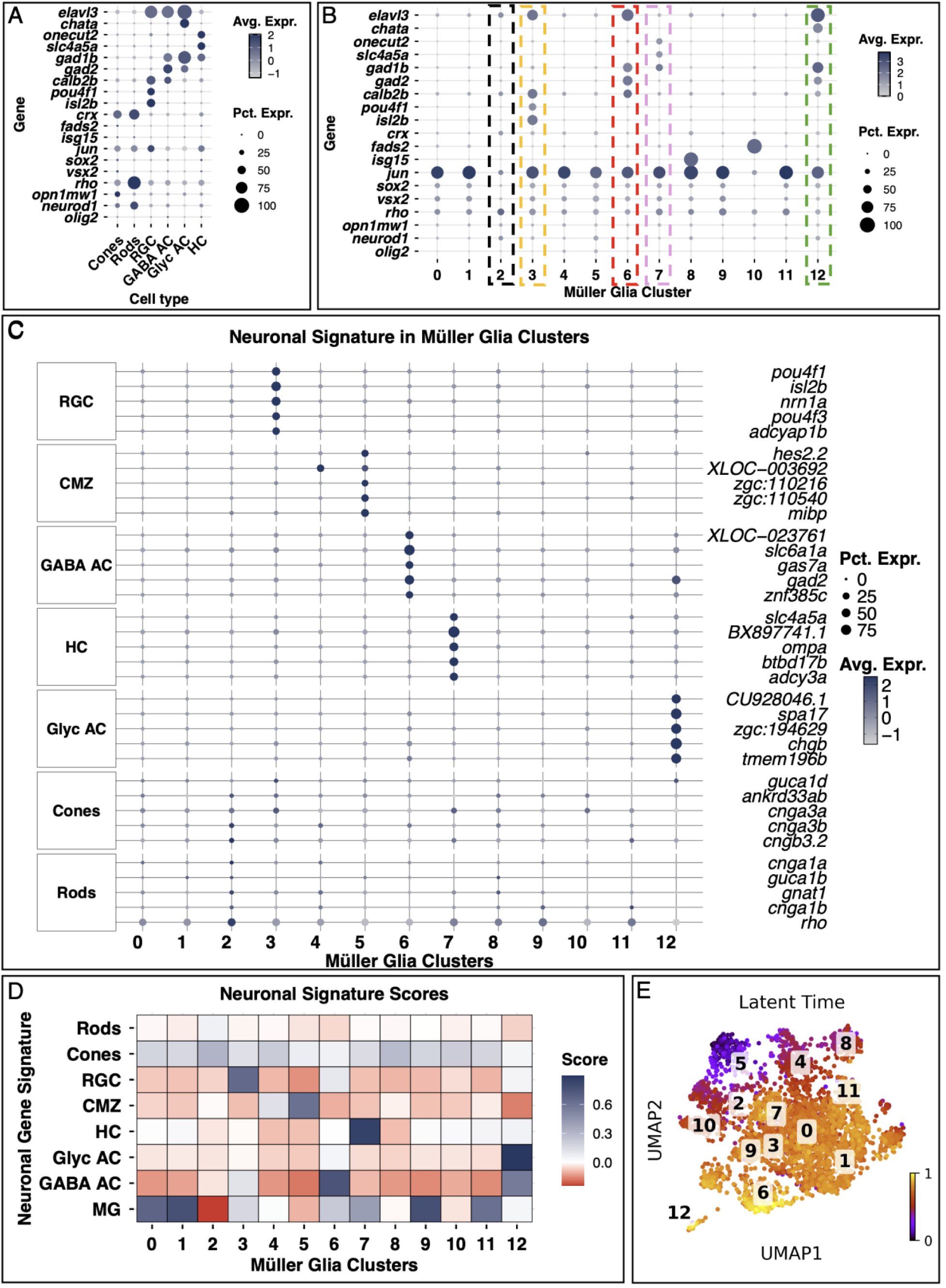
Bioinformatic analysis reveals coherent transcriptional programs in neuron-associated Müller glia clusters. A) Expression dot plot for canonical markers of distinct retinal neuron types in clusters of the 5 dpf zebrafish retinal cell atlas. Abbreviations: RGC = retinal ganglion cells, GABA AC = GABAergic amacrine cells, Glyc AC = glycinergic amacrine cells, HC = horizontal cells. B) Expression dot plot for canonical neuronal markers across clusters within the 5 dpf Müller glia subset. Dashed boxes highlight neuron-associated clusters. C) Expression dot plot for the top 5 differentially enriched genes (DEG) from each retinal cell type projected onto the 5 dpf Müller glia subset. The left facet strip designates retinal cell type source; the right facet strip identifies specific genes. D) Heatmap of scoring output from the Seurat AssignModuleScore function with retinal neurons as a reference (y axis) and Müller glia clusters as a query (x axis). Blue = positive association, red = negative association. E) Latent time analysis demonstrates that neuron-associated Müller glia clusters exhibit similar maturity to that of the core Müller glia.

Unlike the *elavl3*-positive populations, C7 lacked broad pan-neuronal markers, but exhibited a highly horizontal cell specific transcriptional profile, which included the connexin *cx55.5, slc4a5a,* and the transcription factor *onecut2* (Klaassen et al., 2016; Morikawa et al., 2024; Shields et al., 2007) (Figure 2B, pink dashed box). C7 expressed the GABA synthesis enzyme *gad1b* but lacked *gad2*, further distinguishing its metabolic profile from the GABAergic C6 cluster. Notable markers of interneuron progenitors, such as *prox1a* and *ptf1a* (Jusuf & Harris, 2009) showed little to no expression in C7 (data not shown).

C2 showed moderate enrichment for fundamental markers of rod (*rho, saga, sagb, gngt1, gnat1*) and cone (*opn1sw1*) photoreceptor lineages, photoreceptor progenitor identity (*crx*), and the late differentiation stage of bipolar cells (*vsx2*) (Chow et al., 2001; reviewed in Baden et al., 2020) (Figure 2B). This expression profile was reflected in C10. C10 was notable in its enrichment for *fatty acid desaturase 2* (*fads2*), a gene that encodes both Fads1 and Fads2 proteins thought to be neuroprotective against light damage (Ashikawa et al., 2017). Interestingly, C2 and C10 both exhibited a significant downregulation of *jun*, a proto-oncogene transcription factor required for regeneration (Sarich et al., 2025), and *sox2*, a proneural gene involved in maintaining a progenitor state (Pinto & Gotz, 2007; Raymond et al., 2006). This downregulation distinguished C2 and C10 from the proliferating (C5) and immature (C4) populations, yet all four clusters shared reduced expression of the Müller glia core transcriptional signature (Figure 1D), suggesting perhaps a quiescent, lineage-primed state.

To investigate the depth of transcriptional identity shared between Müller glia subtypes and distinct retinal neuron identities, we performed additional bioinformatic comparisons. Our DEG analysis of annotated cell populations within the 5 dpf retinal cell atlas identified sets of cell type specific markers. The compilation of hundreds of cell specific markers yielded a rich transcriptional program for RGCs, glycinergic and GABA-ergic amacrine cells, horizontal cells, and both rod and cone photoreceptors (Supplemental Table 2). In the 5 dpf Müller glia clusters we visualized the expression (by avglog2FC) of top markers from each annotated retinal neuron type and the CMZ to investigate the transcriptional fidelity of enriched non-canonical markers for retinal neuron types and the embryonic-like CMZ, shown by expression dot plot in Figure 2C. This analysis was consistent with our above findings that suggested a prevalence of neuronal markers in distinct Müller glia subtypes. The striking enrichment of an unbiased transcriptional fingerprint suggests a robust association between Müller glia subtypes and distinct neuronal populations.

To explore quantitatively the aggregate expression relationship of specific transcriptional programs within our neuron-associated Müller glia clusters, we used the AddModuleScore function from Seurat. This method calculates a module score for each cell by averaging the expression from the top 50 most enriched genes from each retinal neuron and represents the relative enrichment of a specific neuronal signature for each cluster. These data, shown by heatmap of neuronal signature score in Figure 2D, demonstrated that the genes driving heterogeneity within the subset of neuron-associated Müller glia were distinct from the canonical genes that define glial function. The data also highlighted a striking resemblance of the differentially expressed transcriptome of specific Müller glia clusters to specific neuron-like transcriptional programs. Specifically, C3 exhibited an RGC neuronal signature, C6 and C12 exhibited GABAergic and glycinergic amacrine cell signatures, respectively, and C7 exhibited a horizontal cell signature. To infer the relative maturity of these cells, we performed latent time analysis using scVelo (Bergen et al., 2020). Latent time embeddings projected onto the 5 dpf Müller glia UMAP (Figure 2E) indicated that cells in clusters C3, C6, C7, and C12 shared a similar or advanced maturity with the core, canonical Müller glia populations, distinguishing them from the immature C4/C5 populations. C2, however, appeared to be in an immature state. These unbiased assessments indicate that the neuron-associated clusters are defined by coherent, coordinated neuronal gene programs specific to distinct retinal neuron identities, rather than the sporadic co-expression of isolated transcripts or genes sharing a single common function.

In summary, our analysis reveals that Müller glia clusters C3, C6, C7 and C12 robustly recapitulate neuronal transcriptomic programs specific to RGC, GABAergic amacrine cells, horizontal cells and glycinergic amacrine cells, respectively. Of note, this unbiased analysis also reveals that the proliferative C5 population, and, to a lesser extent, the undifferentiated C4 cohort, exhibit transcriptional programs with marked similarity to those of the peripheral neurogenic CMZ stem cell niche.

### Müller glia *in vivo* express neuron-specific genes and proteins

To validate whether neuronal gene expression occurs in Müller glia *in vivo*, we performed FISH with riboprobes for key markers on transverse sections of the 5 dpf retina alongside Müller glia identification by GS immunostaining. Our scRNAseq data revealed that only a small percentage of Müller glia express such neuron-associated gene markers, and thus we anticipated that these Müller glia phenotypes would be seen infrequently in retinal sections. Indeed, only 295/3563 Müller glia (8.3%) express the C6 gene *gad2* by scRNAseq at 5 dpf (Figure 3A). We identified the occasional *gad2+* Müller glia in the neural retina (Figure 3B,B’; observed in n=6/6 larvae), as seen in an expanded orthogonal view (Figure 3B’) with mRNA expressed diffusely in the cell body. *gad1b* that marked clusters C6, C7 and C12 (Figure 3C; 455/3563 Müller glia, 12.8%) was observed in Müller glia of the neural retina (Figure 3D; n=6/6 larvae), with punctate mRNA ISH signal (Figure 3D’). *onecut2* was specifically expressed in C7 (Figure 3E; 195/3563 Müller glia, 5.5%) with *onecut2* FISH positive puncta in the soma of Müller glia of the neural retina (Figure 3F-F’; n=6/6 larvae). Neuronal transcripts in Müller glia were restricted to the cytosol. The cytosolic localization, combined with the abundance of the same transcripts in adjacent neuronal somata, meant that unambiguous assignment of a FISH signal to a Müller glia required orthogonal re-slicing of the confocal stack on a cell-by-cell basis, making whole-section quantification of these neuron-associated Müller glia intractable by confocal microscopy.

**Figure 3.**
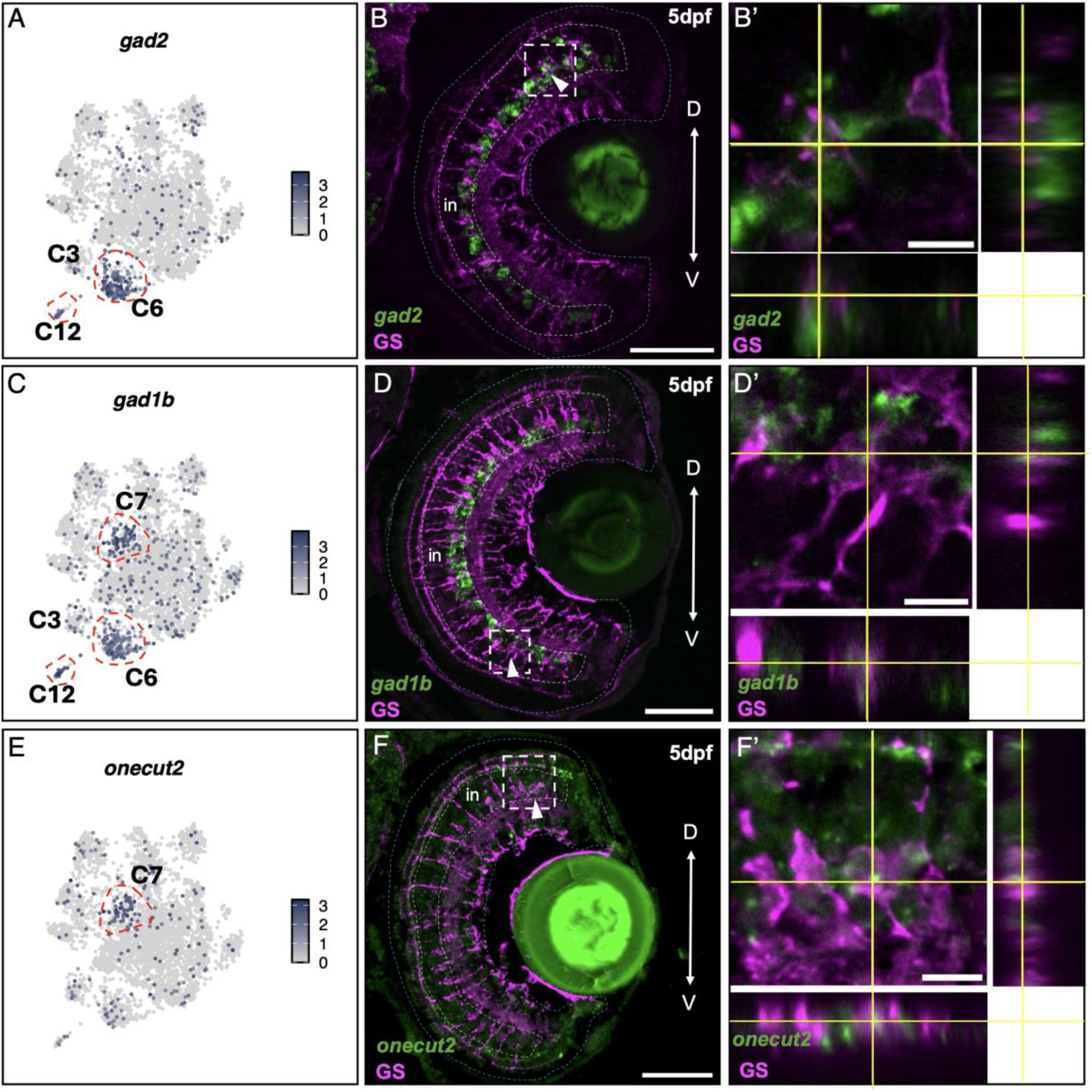
Müller glia subpopulations are defined by canonical transcriptional markers for distinct neuronal populations *in vivo*. A, C, E) UMAP plots of 5 dpf Müller glia scRNAseq data demonstrate the cluster-specific expression of *gad2* (A), *gad1b* (C), and *onecut2* (E). B, D, F) FISH (green) of 5 dpf transverse retinal sections co-labeled with anti-GS immunostaining (magenta). Blue dashed lines delineate eye border. White dashed lines indicate the inner nuclear layer (in). Axes: D = Dorsal, V = Ventral. B’, D’, F’). Enlarged orthogonal views of the regions delineated by white dotted boxes in B, D and F. The y-z (side) and x-z (bottom) dimensions in orthogonal view to demonstrate localization of riboprobe and antibody. White arrowheads (B, D, F) point at cell target for orthogonal view (B’, D’, F’). Scale bars: B, D, F = 50 µm; B’, D’, F’ = 8 µm.

As mRNA does not always equate to protein, we tested for the presence of neuronal-associated proteins in 5 dpf *Tg(gfap:eGFP)* transgenic zebrafish by immunohistochemistry on transverse retinal sections with antibodies that work in fish. Note, we switched to the transgenic fish as the antibodies for GS and Elavl3 were both mouse. We used antibodies for neuron-specific Calbindin expressed by C3 and C6 (Figure 4A; *calb2b* in 666/3563 Müller glia [18.7%] by scRNAseq) and saw striking, but infrequent, expression of Calbindin in the nucleus of Müller glia of the neural retina (Figure 4B-B’; >=1 Calbindin-immunoreactive Müller glia observed in 20/94 retinal sections, n=4/4 larvae). We verified nuclear label of Calbindin in Müller glia through Hoechst label of nuclei (Figure 4C). Notably, Calbindin is found in the nuclei of certain neurons, including rodent cerebellar Purkinje cells and midbrain dopaminergic neurons (German et al., 1997). We visualized the pan-neuronal Elav-family marker with the Hu/CD antibody (Good, 1995) to capture *elavl3-*expressing C3, C6 and C12 Müller glia (Figure 4D; *elavl3* mRNA in 788/3563 Müller glia [22.1%] by scRNAseq). The high density of Hu/CD-immunoreactive tissue in the INL precluded accurate whole-section counting of Hu/CD+ Müller glia by confocal microscopy. However, unambiguous examples of Hu/CD-labelled puncta in the cytosol of eGFP+ Müller glia were confirmed by orthogonal re-slicing (Figure 4E-E’, F; n=7/7 larvae). Together with the FISH data, these findings support that the neuronal transcriptional programs identified in our scRNAseq data result in the production of neuron-specific proteins in discrete Müller glia subpopulations.

**Figure 4.**
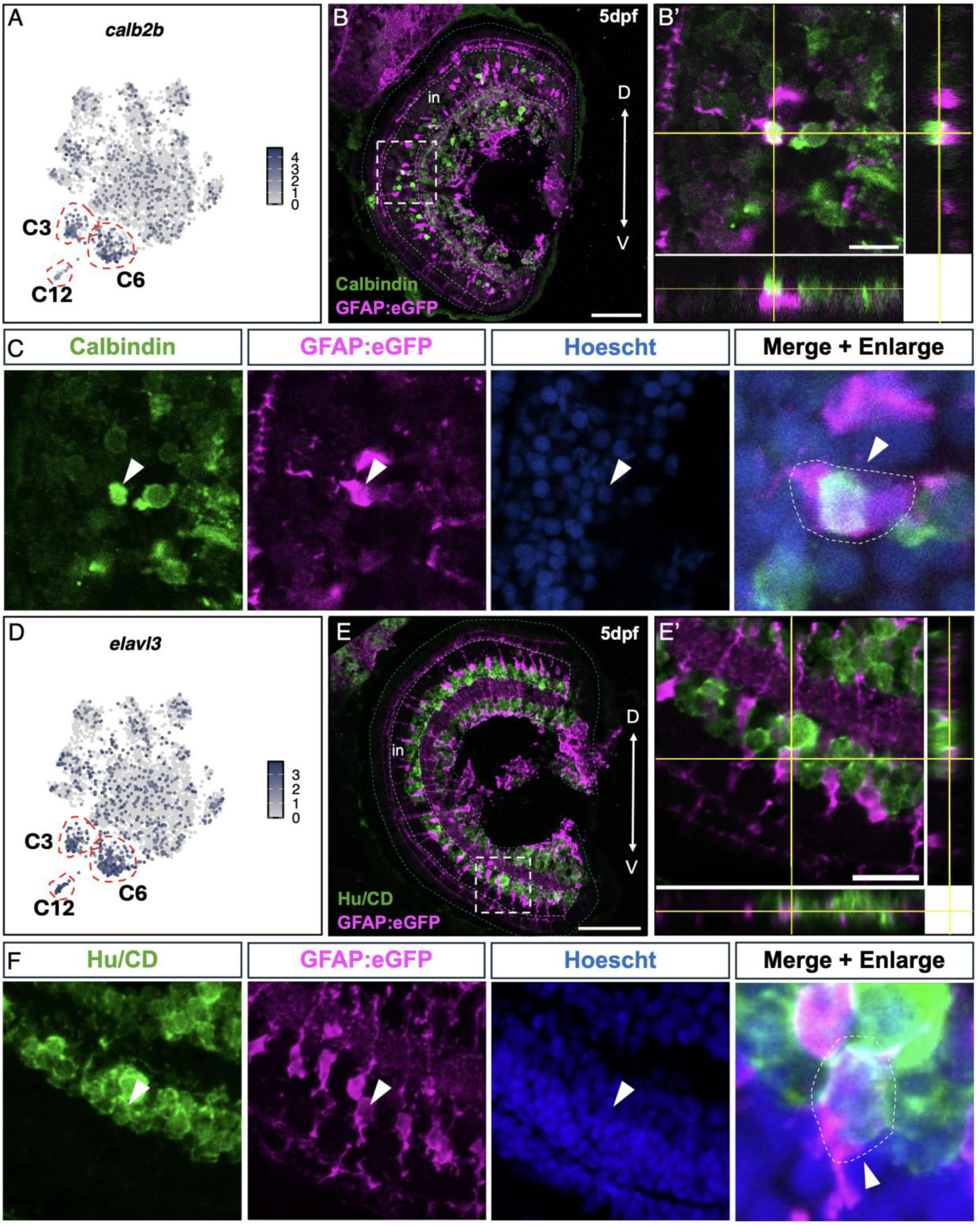
Müller glia subpopulations express neuronal proteins *in vivo*. A,D) UMAP plots demonstrate the expression of *calb2b* (A) and *elavl3* (D) enriched in clusters C3, C6, and C12. B,C,E,F) Immunohistochemistry (green) for Calbindin (B,C) and HuC/D (Elav-family; E,F) on retinal sections of 5 dpf *Tg(gfap:egfp)* zebrafish processed with an anti-GFP JL8 antibody (magenta). Blue dashed lines delineate eye border. White dashed lines indicate the inner nuclear layer (in) (B, E). B’, E’) Enlarged orthogonal views of regions denoted by dashed boxes in B and E. The y-z (side) and x-z (bottom) dimensions in an orthogonal view to demonstrate the subcellular localization of calbindin and HuC/D in eGFP+ Müller glia. Calbindin is expressed in the nucleus of a retinal Müller glia cell (B,B’), and HuC/D in the Müller glia cytoplasm (E,E’), as seen in the enlarged merge with the Hoechst-labeled nuclei (C,F; cell body encircled in white dashes), respectively. White arrowheads represent a specific cell across panels. Inner nuclear layer (INL) shown by blue dashed lines in B. Scale bars: B, E = 50 µm; B’, E’ = 10 µm.

### Spatially distinct Müller glia subpopulations describe a dorso-ventral axis of morphogen patterning

The transcriptomes of clusters C0, C1, C9, and C11 were defined by the highest levels of canonical glial gene expression amongst all clusters and were largely distinct from the transcriptomic signatures of non-glial retinal cell types (Figure 2D). Latent time analysis (scVelo; Figure 2E) indicated that this cohort represented the core mature Müller glia population in the 5 dpf retina. Accordingly, these clusters shared enrichment for fundamental homeostatic markers, including *carbonic anhydrase* (*cahz, ca14*) (Ogilvie et al., 2007), sodium-potassium ATPase subunit *atp1a1b* (Nagashima et al., 2013), *apolipoprotein Eb* (*apoeb*), and prostaglandin metabolism genes (*ptgdsb.1/2*) that become upregulated following light injury in the zebrafish (Sifuentes et al., 2016) (Figure 5A).

**Figure 5.**
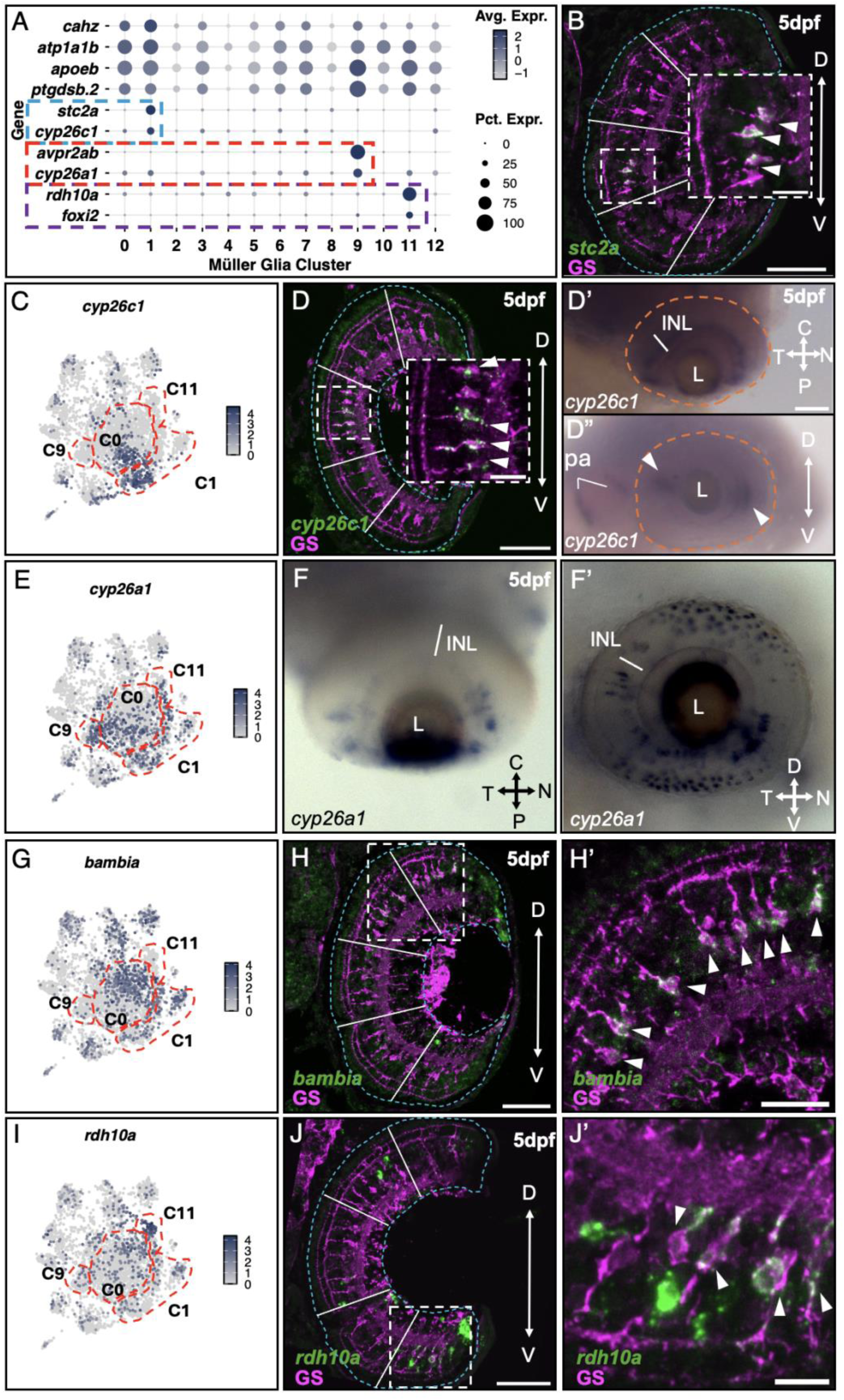
Spatially distinct Müller glia subpopulations suggest a glial-mediated mechanism of dorso-ventral patterning of the retina. A) Dot plot with expression of differentially expressed genes in the 5 dpf Müller glia scRNAseq data. B, C, E, G, I) UMAP plots for *stc2a* (B), *cyp26c1* (C), *cyp26a1* (E), *bambia* (G), and *rdh10a* (I) transcripts within the 5 dpf Müller glia scRNAseq data. Fluorescent *in situ* hybridization (green) on transverse retinal sections co-labeled with anti-GS immunostaining (magenta) to visualize *cyp26c1* (D), *bambia* (H), and *rdh10a* (J). Blue dashed lines delineate eye border. Solid white lines divide the retinal sections into five segments for spatial contextualization: the peripheral retina is defined as both the dorsal and ventral outer segments and includes the CMZ; the central retina spans the three inner segments; the equatorial retina constitutes the centre-most segment. D’, D", F, F’) Whole mount *in situ* hybridization demonstrating the spatial distribution of *cyp26c1* (D’, D") and *cyp26a1* (F, F’). White dashed boxes (B, D, H, J) denote regions enlarged in the corresponding insets (B, D) or adjacent panels (H’, J’). Arrowheads point to riboprobe positive Müller glia. Axes in B, D-D”, F-F’: D = dorsal, V = ventral, C = central, P = peripheral, N = nasal, T = temporal. L = lens. pa = pharyngeal arches. Scale bars: B (inset), D (inset), H’ = 15 µm; B, D, H, J = 50 µm; D’ = 250 µm; J’= 10 µm.

Interestingly, within this mature population we still identified considerable heterogeneity. For instance, C1 was strongly marked by the calcium modulator *stanniocalcin* (*stc2a*) (Boyd & Hyde, 2022; Mitra et al., 2018) and the subfamily C retinoic acid (RA) degradation enzyme *cyp26c1* (Figure 5A, blue dashed box). C9 was distinguished by the *arginine-vasopressin receptor 2a* (*avpr2ab*) and the retinoic acid (RA) degradation enzyme *cyp26a1* (Figure 5A, red dashed box). C11 was characterized by the RA synthesis enzyme *retinol dehydrogenase 10* (*rdh10a*) (Duester, 2008) and the forkhead box transcription factor (*foxi2*), a known ventral marker of the developing zebrafish retina (Solomon et al., 2003) (Figure 5A, purple dashed box). A distinct subset of C0 selectively expressed *bone morphogenetic protein and activin inhibitor homolog a* (*bambia*) (Onichtchouk et al., 1999) (Figure 5G, UMAP expression plot), although these cells were not specifically resolved as an independent cluster by unsupervised clustering.

Our *in vivo* validation at 5 dpf revealed that these molecular signatures map to strikingly distinct spatial domains of the retina. We performed FISH for key markers on transverse retinal sections immunostained with a GS antibody to label Müller glia (Figure 5). We identified a small cluster of *stc2a*+ C1 Müller glia immediately dorsal to the optic nerve head (Figure 5B; expanded inset; observed in 74/74 central retina sections, n=5 larvae). *cyp26c1*+ C1 (and a small C0 subset) Müller glia (Figure 5C; 436/3563 Müller glia, 12.2%) were restricted to a tight band in the central-most retina (Figure 5D; observed in 49/58 central retina sections from n=4 larvae). Whole mount BCIP/NBT ISH for *cyp26c1* as seen in dorsal (Figure 5D’) and lateral (Figure 5D’’, arrowheads) views revealed that expression was restricted to a tight band at the eye equator of consistent thickness that aligned with the visual horizon (n=5 larvae). In contrast, the paralog *cyp26a1* (Figure 5E; 641/3563 Müller glia, 18.0%) was more broadly expressed by Müller glia of the nasal and temporal retina as seen in dorsal (Figure 5F) and lateral views (Figure 5F’) of whole mount ISH (n=5 larvae). Note, however, that this spatial expression pattern was less consistent between animals than the equatorial *cyp26c1* band. *bambia* marked a subset of C0 cells (Figure 5G; 812/3563 Müller glia, 22.8%), and in transverse retinal sections was expressed in the majority of Müller glia of the dorsal peripheral and central retina and not in the ventral retina (Figure 5H,H’; observed in 39/42 central retinal sections, n=6 larvae). Conversely, *rdh10a+* Müller glia (Figure 5I; 520/3563 Müller glia, 14.6%) preferentially populated the ventral peripheral retina (Figure 5J,J’; observed in 54/65 central retinal sections, n=4 larvae), though the occasional *rdh10a*+ cell was observed in the central or dorsal retina (at least one central or dorsal *rdh10a+* Müller glia in 32/65 central retinal sections, n=4 larvae), perhaps reflecting the diverse roles of RA signaling across the tissue. Note, *bambia* was somewhat enriched in C8, which was otherwise explicitly defined by *isg15,* a ubiquitin-like modifier thought to mediate ligase binding activity and immune responses (Langevin et al., 2013). We suspect that C8 (n=135 cells) represents a sparse group of unhealthy cells which made it through the quality control in pre-processing.

### Müller glia subtypes identified at 5 dpf are conserved throughout development into adulthood

To determine if the transcriptional heterogeneity characterized at 5 dpf represents a transient developmental state or a constitutive feature of the zebrafish retina, we integrated our 5 dpf dataset with published scRNAseq data from Müller glia at 9 dpf (Krylov et al., 2023) and the adult (Celotto et al., 2023). This integration and subsequent unbiased clustering yielded a lifespan profile of 14,599 Müller glia organized into 16 distinct integrated clusters. We evaluated integration fidelity for each cluster by Simpson’s Diversity Index (Supplemental Figure S3). After removing two clusters identified as microglia, we annotated seven different clusters based on consistent enrichment of markers for known biology and those identified in our 5 dpf dataset (Figure 6A).

**Figure 6.**
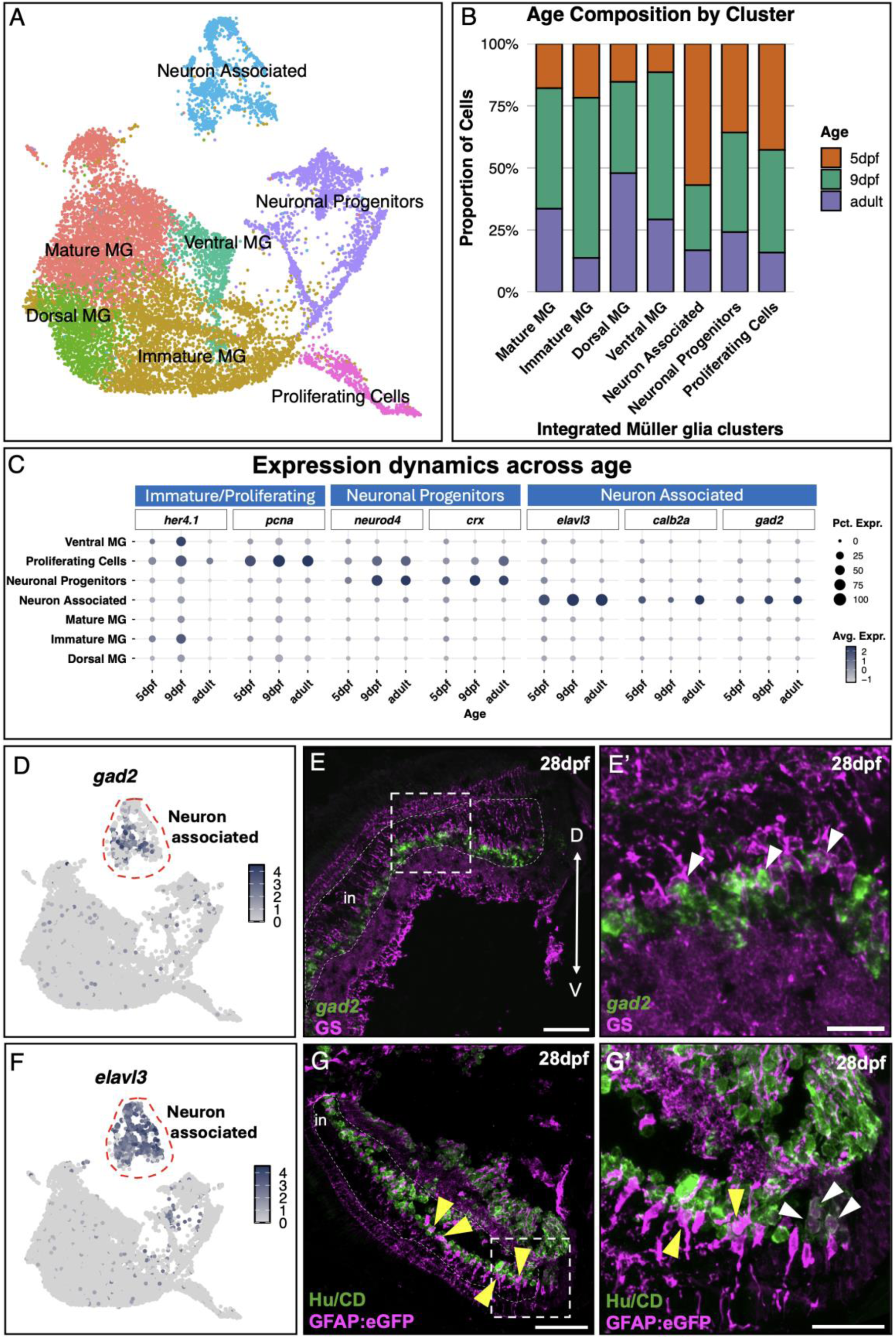
Proliferating, immature, neuron-associated and mature Müller glia subtypes persist from larvae into adulthood. A) UMAP plot of annotated clusters from the integration of 5 dpf, 9 dpf and adult Müller glia scRNAseq datasets. B) Stacked bar chart represents the proportional contribution of each age timepoint to the annotated integrated clusters. C) Dot plot mapping expression of markers of immature/proliferating Müller glia, neuronal progenitors, and neuron-associated Müller glia across distinct clusters (y axis) at each developmental age (x axis) within the integrated dataset. D,F) UMAP of *gad2* (D) and *elavl3* (F) expression in the neuron-associated Müller glia cluster of the integrated scRNAseq data. E,E’) Fluorescent *in situ* hybridization for *gad2* (green) on transverse 28 dpf retinal section immunostained for GS (magenta). The white dashed box (E) indicates the region enlarged in E’. White arrowheads identify *gad2+* Müller glia. Axes: D = dorsal, V = ventral. G-G’) Immunohistochemistry against Hu/CD (green) and eGFP (magenta) on transverse retinal sections from *Tg(gfap:egfp)* 28 dpf fish. The white dashed box in G indicates the region enlarged in G’. White arrowheads identify Hu/CD-labeled presumptive newly born Müller glia. Yellow arrowheads indicate Hu/CD-immunoreactive Müller glia within the central retina. Scale bars: E,G = 50 µm; E’ = 15 µm, G’ = 25 µm.

We calculated the proportion of cells in each of the annotated clusters across the three datasets, as shown by the stacked bar plot in Figure 6B. Each distinct subtype identified at 5 dpf was represented at the later larval and adult timepoints, including: 1) mature Müller glia (canonical Müller glia markers, *apoeb+, aqp1a.1+, cahz+, ptgdsb.2+*); 2) proliferative Müller glia (*pcna+, mki67+, ccnd1+*); 3) Notch-active/immature Müller glia (*her4.1+, her4.2+, her12+*); 4) neuron-associated Müller glia (*elavl3+, pou4f1+, gad2+, vsx1+*); 5) spatially restricted Müller glia defined by morphogen signaling axes (*aldh1a2+, bambia+, cyp26c1+, aldh1a3+, bmpr1ba+*), and 6) Müller glia with a neuronal progenitor identity, distinct from the standard proliferative population, which was revealed with the increased resolution of the integrated dataset. This neuronal progenitor population, present at all ages, was characterized by the strong enrichment for the cone-rod homeobox gene *crx* and markers of neuronal differentiation such as *neurod1* and *neurod4*. The expression data for the immature, proliferating, neuronal progenitor and neuron-associated clusters at each sampled timepoint are shown by dot plot in Figure 6C. All six phenotypes were present at the three ages, though the early larval cells contributed more of the neuron-associated Müller glia and fewer of the dorsal and ventral Müller glia.

To confirm that the lifespan persistence of these subtypes is biological rather than a technical artifact of integration, we performed FISH on transverse retinal sections at 28 dpf. We examined the expression of the same neuron-associated markers (*gad2, gad1b*) we observed at 5 dpf by FISH (Figure 3B, B’, D, D’) at 28 dpf on retinal sections immunostained for GS. We identified expression of *gad2* in individual Müller glia (Figure 6D; expressed in 199/4021 [4.9%] adult Müller glia by scRNAseq) at 28 dpf (Figure 6E-E’; observed in n=4 juveniles). Further, as *elavl3* defined neuron-associated clusters (Figure 6F; *elavl3* mRNA expressed in 270/4021 [6.7%] adult Müller glia by scRNAseq), we performed immunohistochemistry for Elav protein on retinal sections from 28 dpf *Tg(gfap:egfp)* fish. We identified double positive Hu/CD+/eGFP+ in Müller glia of both the central retina (Figure 6G-G’; yellow arrowheads; observed in n=4 juveniles) and the retinal periphery adjacent to the CMZ (Figure 6G’; white arrowheads; observed in n=4 juveniles). At 28 dpf the *gfap* promoter-driven eGFP signal attenuated with distance from the retinal periphery, thus we did not characterize fully the extent of HuC/D immunoreactivity in the most central Müller glia. These data confirm that neuron-associated glial subtypes are not a transient embryonic phenomenon but a constitutive feature of the zebrafish retina that persists into early adulthood.

### Müller glia subtypes that define spatially distributed BMP and RA signaling axes remain into adulthood

Our 5 dpf analysis revealed a nascent dorso-ventral axis of Müller glia defined by their expression of players in RA synthesis and BMP signaling (*rdh10a+, bambia+*), bisected by an equatorial band of *cyp26c1+/stc2a+* Müller glia. To determine if this spatial organization is a constitutive feature of the zebrafish retina, we examined expression patterns within our integrated 5 dpf, 9 dpf, and adult Müller glia scRNAseq dataset. Transcriptional profiling across the lifespan revealed that these spatially distinct subtypes persisted into adulthood (Figures 6B, 7A). Leveraging the spatial markers identified at 5 dpf within the integrated dataset allowed us to resolve additional dorso-ventral patterning genes as marking spatially distinct Müller glia. These included mRNAs for: 1) the canonical dorsal transcription factors Tbx5a*/*b (Pi-Roig et al., 2014), 2) BMP pathway components, specifically the ligands *growth and differentiation factor 6a* (*gdf6a*) and Bmp4 (French et al., 2009; Gosse & Baier, 2009) in dorsal Müller glia, and the BMP receptor *bmpr1ba* (Liu et al., 2003)in ventral Müller glia, and 3) the RA signaling component *dhrs3a* in ventral Müller glia.

Interestingly, we observed a temporal shift in the contribution of specific regulators of the RA morphogen in our bioinformatic analysis. We found that while the upstream RA synthesis enzyme *rdh10a* and *cyp26c1* defined the axis in both early larvae and adult fish, missing at 5 dpf were the downstream RA synthesis enzymes *aldh1a2* (dorsal) and *aldh1a3* (ventral), which specify dorsal and ventral domains during eye field development (Duester, 2022). The mRNAs for these two enzymes appear to be progressively upregulated by dorsal and ventral Müller glia as the animal matures. These findings place strong emphasis on the importance of the RA axis in the post-embryonic retina.

We validated this developmental trajectory *in vivo* via FISH on transverse retinal sections processed for GS immunostaining to identify Müller glia (Figure 7). First, we examined the RA synthesis domains. *aldh1a2* and *aldh1a3* expression was virtually absent in Müller glia of the 5 dpf retina (*aldh1a2* expressed in 17/3563 Müller glia [0.5%] at 5 dpf; *aldh1a3* expressed in 89/3563 Müller glia [2.5%] at 5 dpf), though at this timepoint robust expression was observed for both genes in dorsal (*aldh1a2*) and dorsal/ventral (*aldh1a3*) CMZ (data not shown). Consistent with our bioinformatic data (*aldh1a2* expressed in 124/7015 Müller glia [1.8%] at 9 dpf by scRNAseq; *aldh1a3* expressed in 514/7015 Müller glia [7.3%] at 9 dpf by scRNAseq), *aldh1a2* (dorsal) and *aldh1a3* (ventral) mRNAs emerged *in vivo* in peripheral Müller glia at 9 dpf (Figure 7B, D; dorsal *aldh1a2* bias observed in 108/122 retinal sections, n=11 larvae; ventral *aldh1a3* bias observed in 152/153 retinal sections, n=5 larvae). The spatially biased expression of RA synthesis enzymes in Müller glia was maintained at 28 dpf (Figure 7C-C’, E-E’; dorsal *aldh1a2* bias observed in 150/153 retinal sections, n=4 juveniles; ventral *aldh1a3* bias observed in 99/104 retinal sections, n=4 juveniles), and remains consistent with our bioinformatic data (*aldh1a2* expressed in 786/4021 adult Müller glia [19.5%] by scRNAseq; *aldh1a3* expressed in 937/4021 adult Müller glia [23.3%]); Müller glia expression outside of the defined regions remained extremely limited, making co-expression challenging to quantify. Both *aldh1a2* and *aldh1a3* were expressed by both the dorsal and the ventral CMZ at 9 dpf through 28 dpf. These expression data argue that the glial-associated RA synthesis domains mature as larval development proceeds.

**Figure 7.**
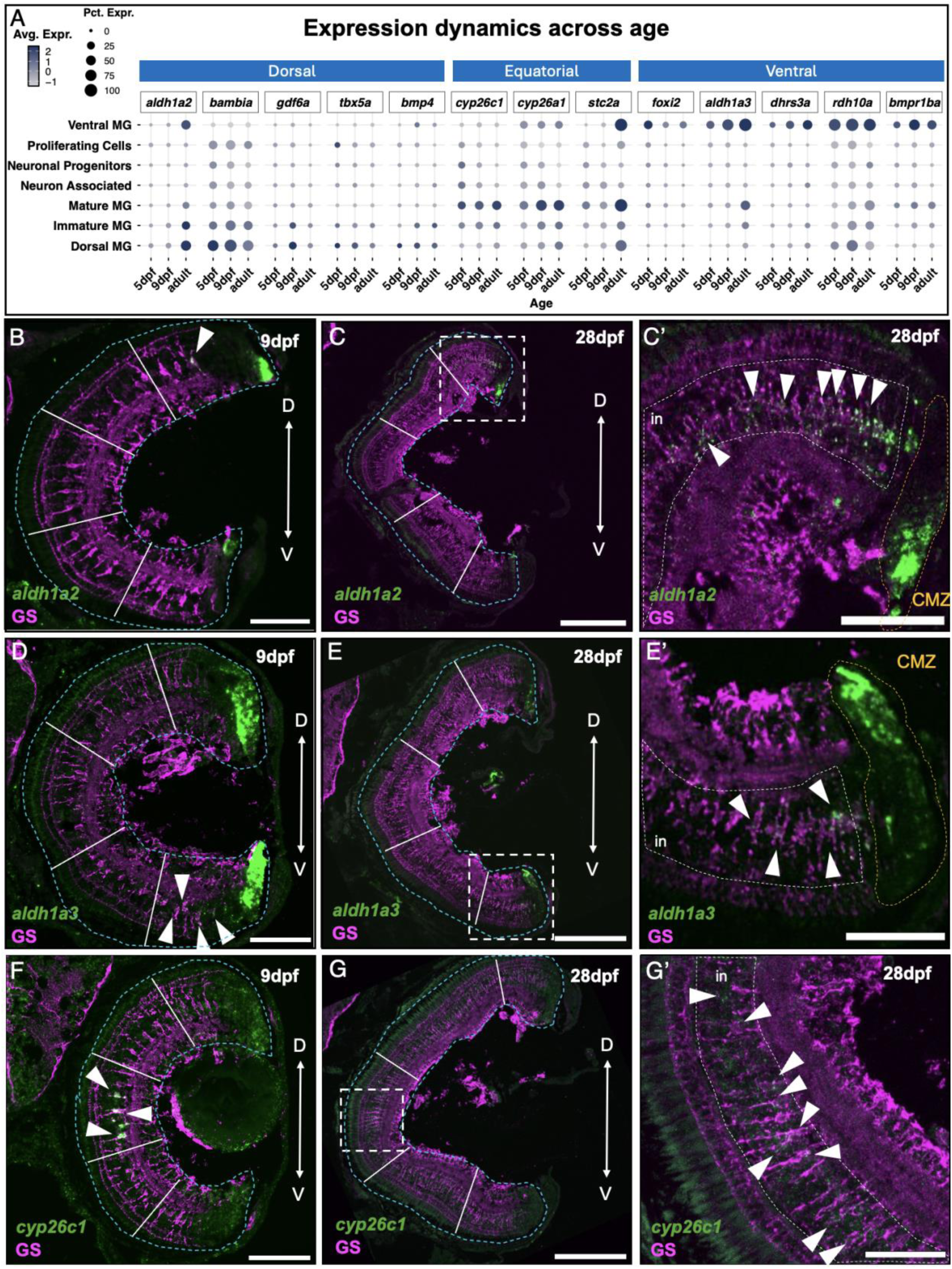
Müller glia subpopulations define a dorso-ventral axis of RA metabolism across the lifespan of the zebrafish. A) Expression dot plot for markers of dorsal, equatorial and ventral Müller glia across distinct clusters (y axis) for each developmental age (x axis) in the integrated dataset. B-G’) Fluorescent *in situ* hybridization (green) on transverse retinal sections of 9 dpf (B, D, F) and 28 dpf (C, E, G) zebrafish co-labeled for GS immunostaining (magenta). Blue dashed lines delineate eye border. Solid white lines divide the retinal sections into five segments for spatial contextualization: the peripheral retina is defined as both the dorsal and ventral outer segments and includes the CMZ; the central retina spans the three inner segments; the equatorial retina constitutes the centre-most segment. White dashed lines indicate the inner nuclear layer (in). Riboprobes target *aldh1a2* (B-C’), *aldh1a3* (D-E’), and *cyp26c1* (F-G’). Dashed boxes (C, E, G) indicate the regions enlarged in the corresponding adjacent panels (C’, E’, G). White arrowheads identify riboprobe positive Müller glia. Ciliary marginal zone is shown by orange dashed lines at 28 dpf in C’ and E’. Axes: D = dorsal, V = ventral. Scale bars: B, D, F = 50 µm; C, E, G = 150 µm; C’, E’ and G’ = 50 µm.

The RA degradation enzyme *cyp26c1* exhibited a distinct developmental trajectory. The precise equatorial *cyp26c1* mRNA expression identified at 5 dpf (Figure 5D) and 9 dpf (Figure 7F) expanded and became distributed throughout the central retina by 28 dpf (Figure 7G,G’). Our bioinformatics data reveal that the expression of *cyp26c1* persisted at a comparable proportion across development: 14.9% at 5 dpf, 16.6% at 9 dpf, and 16.9% in the adult. Collectively, these data reveal an increase in mRNAs of the molecular machinery for RA synthesis in Müller glia of the peripheral retina as the organism matures, while the degradation element in Müller glia of the central retina appears to remain proportional and spatially consistent.

### Neuron-associated Müller glia subtypes are evolutionarily conserved, while spatial signaling axes are likely specific to the zebrafish

To determine if Müller glia transcriptional heterogeneity represents a fundamental feature of vertebrate retinas or a specialization of the teleost lineage, we compared our integrated zebrafish Müller glia atlas with publicly available datasets from the embryonic chicken (Yamagata et al., 2021), mouse (Li et al., 2024), fetal human (Kriukov et al., 2025; Lu et al., 2020; Sridhar et al., 2020), and adult human (Li et al., 2026) retinas.

Briefly, processed chick Müller glia data were obtained from the Broad Single Cell Portal (Figure 8A, n=3341 cells). Müller glia from the mouse (Figure 8B, n=8204 cells), fetal human (Figure 8C, n=7133 cells) and adult human (Figure 8D, n=47,235 cells) were bioinformatically isolated from retinal atlases obtained from CELLXGENE (CZI Single-Cell Biology et al., 2023) and processed as described in Methods. For consistency, zebrafish gene nomenclature is used hereafter to refer to orthologs in all species.

**Figure 8.**
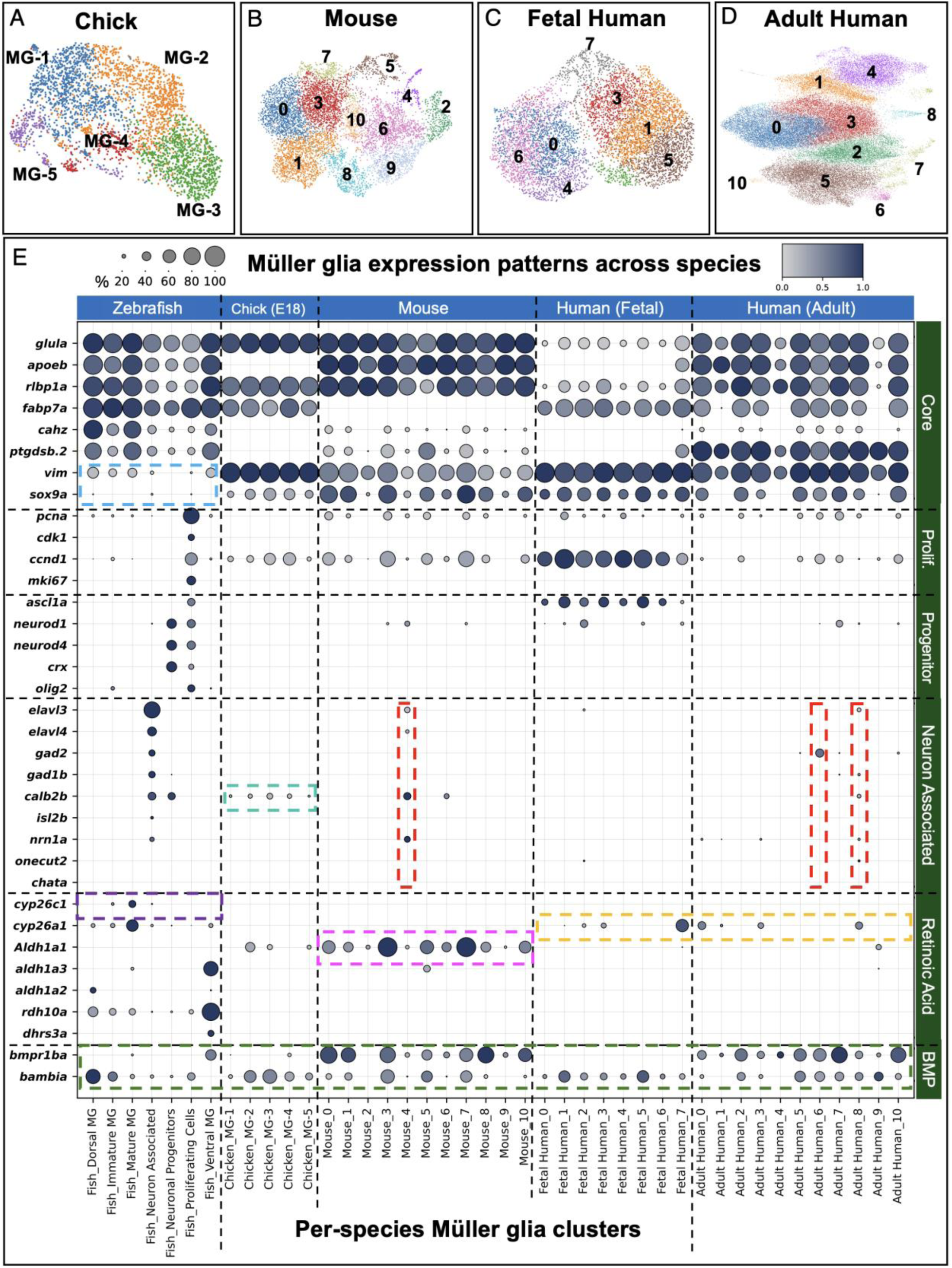
Müller glia subtypes identified in the zebrafish are conserved in the chicken, mouse and human. A-D) UMAP plots of Müller glia publicly available scRNAseq data are shown for the E18 chick (A), post-embryonic mouse (B), fetal human (C), and adult human (D). E) Expression dot plot for the integrated zebrafish dataset and each of the species mentioned above. Blue column headers denote species/dataset. The bottom x-axis identifies clusters corresponding to panels A-D. The right y-axis (green headers) indicates functional grouping; the left y-axis lists individual genes. Coloured dashed boxes referenced in text.

First, we examined core glial identity, based on markers we identified in zebrafish (Figure 1D). While we observed broad conservation of metabolic markers such as *glula* and *rlbp1a* (zebrafish, chicken, mouse, adult human), we noted species-specific variances in lipid carriers (*apoeb* absent in chicken; *fabp7a* absent in mouse) and prostaglandin metabolism (*ptgdsb.2* absent in chicken) (Figure 8E). Classical mammalian markers like *vimentin* (*vim*) and *sox9* were robustly expressed in chick, mouse, and human, but were absent or expressed at low levels in zebrafish Müller glia (Figure 8E, blue dashed box).

We investigated the conservation of the novel subpopulations identified in our zebrafish atlas. Given the limited regenerative capacity of mammals, we were surprised to observe markers of active proliferation (*pcna*) and cell-cycle (*cdk1*, *ccnd1*, *mki67*) broadly expressed at low levels in the adult mammalian datasets. Progenitor identity markers were largely absent in the adult mouse and human datasets, though *ascl1a* and *ccnd1* were robustly expressed in human fetal glia. We did, however, identify discrete subsets of mammalian Müller glia that recapitulated the neuron-associated phenotypes observed in zebrafish. Specifically, we identified a cluster in the mouse (C4) characterized by expression of *calb2b*, *nrn1a* and *elavl3/4,* and two subsets of adult human Müller glia enriched for the GABAergic marker *gad2* (C6) and *elavl3, gad1b, calb2b, onecut2* and *nrn1a* (C8) (Figure 8E, red dashed boxes). We saw weak expression of *calb2b* marker in all clusters in the chick (Figure 8E, teal dashed box).

These data suggest that the capacity for Müller glia to maintain neuron-specific transcriptional programs is an evolutionarily conserved feature of the vertebrate retina, present even in species that lack robust regenerative potential.

Finally, we examined the key players of the zebrafish spatial morphogen axes. While transcripts for *aldh1a2* and *aldh1a3* RA synthesis enzymes were virtually absent in the chicken, mouse and human, the paralog *Aldh1a1* was robustly expressed in Müller glia of the postnatal mouse, particularly in clusters C3 and C7 (Figure 8E, magenta dashed box). Of note, *aldh1a1* does not exist in the zebrafish genome (Pittlik et al., 2008). The equatorial RA degradation marker *cyp26c1,* which was strongly expressed in Müller glia of the larval zebrafish retinal equator, was specific to the teleost lineage (Figure 8E, purple dashed box). The *cyp26a1* paralog was expressed in human Müller glia at both the fetal and adult stages (Figure 8E, orange dashed box), however, the mouse and chick Müller glia lacked expression of either *cyp26* gene. Furthermore, *bambia,* which defined the dorsal Müller glia in the zebrafish, was expressed by most Müller glia clusters in the chicken, mouse and human. Similarly, the BMP receptor *bmpr1ba,* which defined ventral Müller glia in zebrafish, was expressed by most clusters of the mouse and adult human but was absent in the chick and fetal human (Figure 8E, green dashed box). These data suggest a broad role for BMP signaling in postnatal mammalian Müller glia.

## Discussion

We find that Müller glia heterogeneity in the zebrafish is established as early as 5 dpf, characterized by transcriptionally and spatially distinct subtypes. While previous work suggested heterogeneity in quiescent Müller glia at later larval stages (Krylov et al., 2023) and in adults (Celotto et al., 2023; H. Xu et al., 2025), our data demonstrate that Müller glia diversity is encoded much earlier, concurrent with the maturation of retinal circuitry (Wan et al., 2016). We identify by bioinformatics and *in vivo* validation three major populations that present at an early larval stage and persist into adulthood: 1) a proliferative and immature population in both the peripheral and central retina; 2) a novel cohort of neuron-associated Müller glia that express coherent transcriptional programs specific to distinct neuronal subtypes, including retinal ganglion, amacrine and horizontal cells; and 3) spatially distinct mature Müller glia subsets that define a dorso-ventral axis of retinoic acid metabolism, bisected by a novel cyp26c1-expressing equatorial domain. Furthermore, our cross-species comparison suggests that both neuron-associated gene programs and the capacity for broad postnatal morphogen metabolism are evolutionarily conserved traits shared by mammalian Müller glia. These findings, summarized by schematic in Figure 9, advance our understanding of Müller glia beyond their established roles, reframing them as functionally heterogeneous populations and introducing potential novel roles, such as establishing distinct retinal microenvironments.

**Figure 9.**
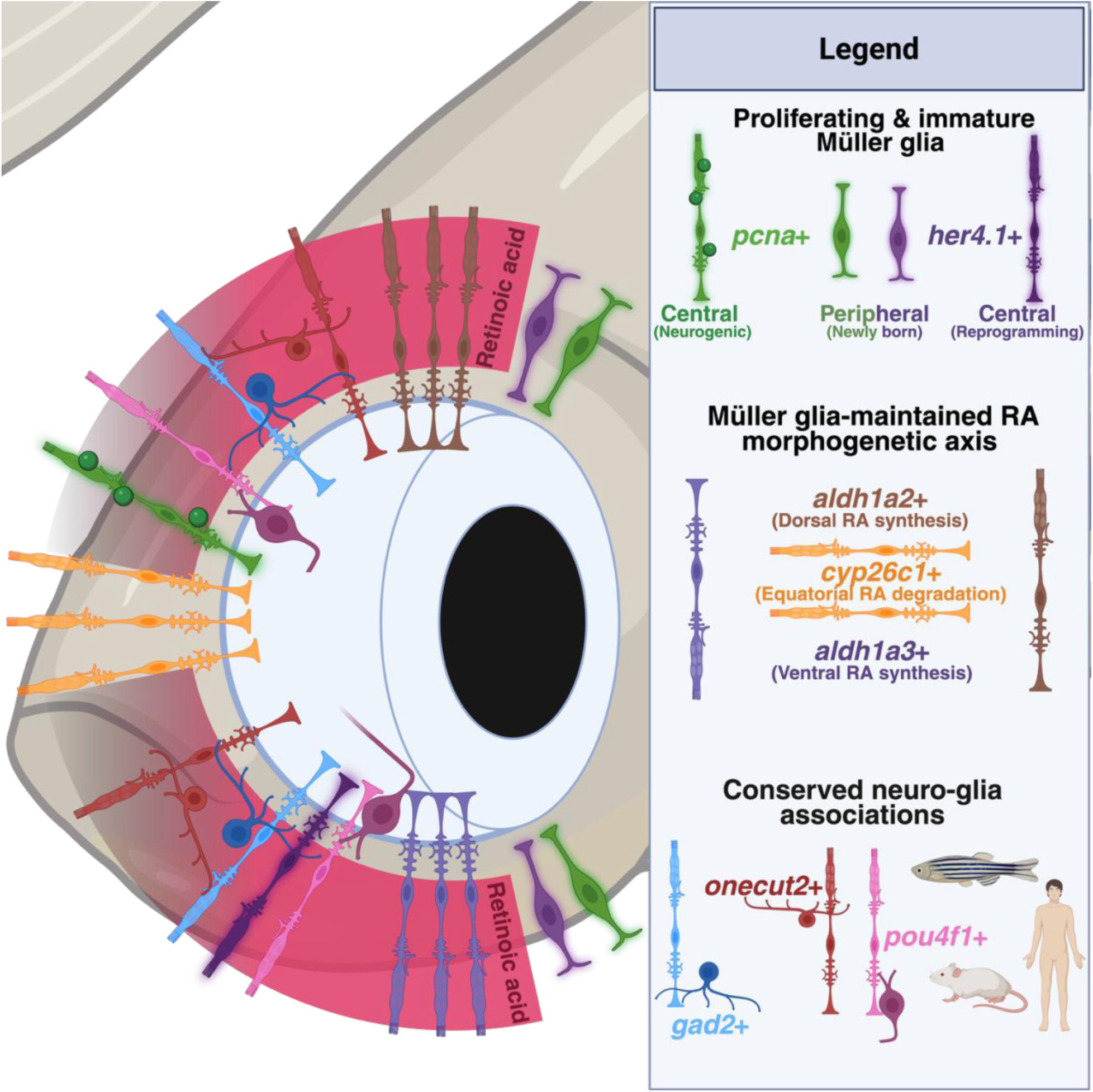
Schematic summary of Müller glia subtypes identified throughout zebrafish across development and conserved between species. Three constitutive Müller glia subpopulations persist from larval stages into adulthood: a proliferative, immature population in the peripheral and central retina; neuron-associated Müller glia expressing coherent transcriptional programs matched to distinct neuronal subtypes (retinal ganglion, amacrine, and horizontal cells); and spatially distinct subsets that define a dorso-ventral axis of retinoic acid metabolism, bisected by a *cyp26c1*-expressing equatorial domain. Cross-species comparison indicates that neuron-associated programs in Müller glia appear evolutionarily conserved in mammals, while the morphogen-metabolizing axis established by Müller glia may exhibit teleost-specific specializations.

A central finding of this work is the identification of Müller glia subpopulations that express coherent distinct neuronal gene programs alongside their core glial identity. Our analysis reveals that this is not the sparse, coincidental expression of a few isolated markers, but rather the coordinated co-expression of transcriptional programs differentially enriched in distinct Müller glia subtypes to match distinct neuronal counterparts that include retinal ganglion, glycinergic or GABAergic amacrine, and horizontal cells. This discovery significantly expands upon existing literature, which has primarily documented neuron-associated transcriptional profiles within the context of injury-induced reprogramming (Celotto et al., 2023; Krylov et al., 2023; H. Xu et al., 2025). Our data demonstrate that this transcriptional diversity is not just a reactive state but a constitutive feature of the uninjured retina, providing evidence for intrinsic functional heterogeneity in zebrafish Müller glia.

One possibility is that the neuron-associated Müller glia are an artifact of the single-cell isolation process. Müller glia extend highly ramified processes that intimately ensheathe virtually all retinal neurons (Lenkowski & Raymond, 2014; Wang et al., 2017). Thin neuronal protrusions or membrane remnants could escape the enzymatic and mechanical tissue dissociation and remain stuck to glial cells, potentially introducing exogenous transcripts during the single-cell capture. For several reasons, however, we argue this possibility is unlikely. First, the neuron-associated transcriptomes were comprehensive, consisting of multiple mRNAs encoding proteins with various functions, including transcription factors whose mRNA we would not expect to be found in neuronal processes. Second, Müller glia showed no enrichment for predicted doublets: Scrublet (Wolock et al., 2019) flagged only 3.6% of Müller glia as doublets, placing them mid-range across retinal cell types and below populations such as corneal fibroblasts (8.7%) and glycinergic amacrine cells (7.2%), arguing against their arising from co-captured glia–neuron pairs. Third, while contamination from ambient RNA is a real phenomenon, one would expect ambient mRNAs to be stochastic in their distribution, rather than aggregate within specific cells to form segregated and neuron-type matched programs in specific Müller glia subtypes. Finally, our independent *in vivo* validation using FISH and immunohistochemistry did identify individual Müller glia that appeared to express neuronal markers. Most convincingly, we identified nuclear Calbindin immunoreactivity localized intracellularly within Müller glia cell bodies in intact tissue (Figure 4C), a spatial distribution that cannot be explained by external membrane adhesion from dissociated neurons. Two potential approaches could in the future provide further validation of these populations. First, definitively establishing that these neuronal transcripts are produced by Müller glia, rather than adhering to them, would benefit from single-nucleus RNA sequencing. Second, enumerating these populations defined by cytosolic markers *in situ* for a comprehensive spatial quantification will require future exploration with more sensitive tools such as RNAscope, expansion microscopy or super-resolution microscopy.

Strikingly, our cross-species analysis revealed that neuron-associated gene expression identified in zebrafish Müller glia is conserved in higher vertebrates. We identified populations of *calb2b+* Müller glia in the chicken, mouse, and human, alongside broader neuronal programs in distinct clusters within both mouse and adult human datasets. This evolutionary conservation suggests that neuron-associated heterogeneity is a fundamental feature of vertebrate retinal physiology, representing a functional specialization where glial subtypes are transcriptionally tuned to support discrete neuronal populations. Recent work by Brown et al. (2025) demonstrates such neuro-glial coupling in the mouse, showing that developing Müller glia intimately associate with and functionally modulate intrinsically photosensitive RGCs to shape circadian circuits. Our data suggest this tailored support extends to other retinal neurons; for example, the expression of GABA synthesis and transport machinery (*gad1b, gad2, slc6a1a*) in C6 Müller glia could facilitate localized neurotransmitter recycling for nearby GABAergic amacrine cells.

However, in the context of the regenerative zebrafish retina, we must consider an alternative biological role. Our integrated lifespan analysis in the zebrafish reveals these neuron-associated populations persist across developmental time. While we did not observe the expression of proneural genes in neuron-associated clusters during homeostasis, the persistence of neuron-associated populations raises the possibility that they function as a quiescent, lineage-restricted progenitor pool. Possibly, Müller glia feature an enhanced neurogenic competency in larval zebrafish in response to the increased demands for neurogenic volume and diversity during the dramatic eye growth of early retinal development. This broadened developmental potency may subsequently restrict to predominantly rod genesis in the mature adult retina, a hypothesis that requires future validation via *in vivo* lineage tracing.

The canonical Müller glia neurogenesis pathway involves the generation of multipotent progenitors that predominantly produce rod photoreceptors (Bernardos et al., 2007; Kustermann et al., 2010). We did not observe a discrete ‘rod-associated’ population; instead, we found pan-glial expression of the rod opsin *rho* and strong expression of *crx* in the neuronal progenitor cluster. This expression is consistent with the established model that all zebrafish Müller glia possess the latent capacity to generate rod photoreceptors. Conversely, if Müller glia are also the source of amacrine or RGC neuronal turnover under homeostatic conditions, they would likely require a distinct, specialized starting state. Thus, these neuron-associated clusters may serve a dual role: providing specialized metabolic support to their neuronal counterparts during homeostasis, while retaining a lineage-biased potential to efficiently regenerate specific neuronal subtypes upon injury.

We also identified a distinct set of Müller glia subtypes that we hypothesize act as key regulatory nodes in retinal spatial patterning. We describe what appears to be a robust dorso-ventral axis of RA metabolism maintained by Müller glia. While RA gradients are known to specify dorsal/ventral identity in the developing eye field (Duester, 2013, 2022), these data suggest the necessity for this gradient in the differentiated, mature retina. Alongside the dorso-ventral axis of RA synthesis, we identify a distinctive Müller glia subpopulation expressing the degradation enzyme *cyp26c1* in a tight equatorial band at 5 dpf and 9 dpf. This band effectively bisects the dorsal (*aldh1a2*) and ventral (*aldh1a3*) synthesis domains. These data argue for a glial-maintained RA gradient that is sharpened by RA degradation in the maturing larval retina (5-9 dpf), when the distance from the RA source at the retinal periphery to the equator is relatively short. The organization of *cyp26c1* expression in the zebrafish retina mirrors the wedge and bullseye patterning of *Cyp26a1/c1* observed in the chick and mouse retinas (Da Silva & Cepko, 2017; McCaffery et al., 1999; Sakai et al., 2004). Our findings support the idea of an evolutionarily conserved mechanism of RA degradation at the zebrafish retinal equator. However, our cross-species analysis reveals that *CYP26C1/Cyp26c1* are not expressed by Müller glia in the embryonic chick or postnatal mouse. This absence suggests that while the equatorial ‘wedge’ pattern is conserved, the transcriptional responsibility for RA catabolism in the zebrafish retina is uniquely shifted to specialized *cyp26c1*-expressing Müller glia.

The spatial restriction of RA-catabolizing Müller glia likely contributes to localized photoreceptor patterning. Consistently, Cyp26 enzymes establish high-acuity zones in chicks and humans (Da Silva & Cepko, 2017), while localized glial *cyp26* expression drives UV cone outer segment elongation in the zebrafish acute zone (Lahne et al., 2026). Furthermore, the equatorial *cyp26c1*+ glial band in larval zebrafish closely mirrors high-density *opn1sw2*+ (blue) cone domains and the rod-free zone (Zimmermann et al., 2018) . Consequently, these equatorial glia may regulate local RA availability to shape high-acuity cone opsin gradients and cell fate patterning, a hypothesis requiring future functional validation.

Importantly, the RA morphogen axis is not a transient phenomenon. At later ages (28 dpf), when the eye is much larger, *aldh1a2* and *aldh1a3* expression becomes enriched in Müller glia of the dorsal and ventral retinal periphery, where cells transition from genesis at the CMZ to their mature fate. Similarly, the regional domain for RA degradation at the central eye found in larval retina appears to expand from a tightly regulated band, four or five *cyp26c1+* Müller glia in width, to diffuse *cyp26c1* expression in both Müller glia and other cell types of central retina at 28 dpf. In the zebrafish, the retina grows continuously throughout life, adding new neurons at the ciliary margin that must be integrated into existing circuits (Karl & Reh, 2010; reviewed in Hoon et al., 2014; reviewed in Lahne et al., 2020). In this respect, the adult persistence of the morphogen-synthesizing axis and the spatial relationship of both RA synthesis and BMP signaling domains to the dorsal and ventral peripheral retina adjacent to the CMZ may have interesting implications for the regulation of CMZ-mediated neurogenesis and/or gliogenesis (Ueki et al., 2015). Further, RA signaling is a potent regulator of stemness, capable of antagonizing fibroblast growth factor (Fgf) signaling to restrict progenitor proliferation (Duester, 2013; Todd et al., 2018). It is therefore noteworthy that an *fgf24*-expressing central Müller glia population described by Krylov et al. (2023) is bordered precisely by the RA synthesis domains we describe here.

Future work is required to determine whether the spatially restricted organization of RA-associated Müller glia is restricted to the teleost lineage. While we found mice express *Aldh1a1* in Müller glia and lack *cyp26c1*, analysis of protein or mRNA within the mouse retina is needed to determine if these mammalian transcripts exhibit spatial bias. We hypothesize, however, that the spatial compartmentalization observed in zebrafish reflects a divergence in growth strategies. This divergence also highlights the distinct transcriptomic shift we observed between the fetal and adult human Müller glia. In the human datasets, progenitor-associated markers (*ascl1a, ccnd1*) were robustly expressed in fetal tissue but lost to adulthood, reflecting a finite and strict temporal development window. In contrast, the zebrafish sustains these immature transcriptomic signatures and spatial morphogen gradients across its lifespan to facilitate ongoing neurogenesis at the ciliary margin. Therefore, while the static mammalian retina undergoes terminal maturation, the continuously growing zebrafish retina may require a permanent, glial-maintained morphogen scaffold to instruct lifelong retinal cell differentiation.

## Conclusion

We present a comprehensive single-cell atlas of the developing zebrafish retina and provide foundational evidence that vertebrate Müller glia feature a diversity of spatially and functionally distinct subtypes. Further, by characterizing these populations through the lifespan of the regenerative zebrafish and revealing the evolutionary conservation of defining transcriptional programs in the chick, mouse and human, our findings set a translational framework for novel exploration of glial heterogeneity in the mammalian retina.

## Acknowledgements

The authors would like to thank Dr. Sarah Childs for the use of her fish facility, Drs. Jeff Biernaskie and Nicole Rosin for performing single-cell RNA sequencing and bioinformatic alignment, and ITSR Labs Inc. (Calgary, Canada) for use of their computational resources. S.S.S. was supported by the Alberta Graduate Excellence Scholarship (AGES), Spinal Cord, Nerve Injury and Pain (SCNIP) Graduate Award from the Hotchkiss Brain Institute and the Cumming School of Medicine Graduate Scholarship. This work was supported by a project grant from the Canadian Institutes for Health Research (PJT-178044), support from the Roy and Joan Allen Chair in Vision and Visual Sciences to S.M., support from the Alberta Ride for Sight and an award from the Lions Sight Centre Fund to S.M.

## Ethics approval Statement

All animal procedures were performed in accordance with the guidelines established by the Canadian Council on Animal Care (CCAC) and were approved by the University of Calgary Animal Care Committee.

## Data Availability Statement

The data and code that support the findings of this study are openly available. The code used for data processing, analysis and figure generation can be found in the online repository https://github.com/s-storey/storey-muller-glia-heterogeneity-2026.

Processed analysis datasets for the datasets generated in this study and those used for analysis and figure generation can be accessed by the Zenodo repository at 10.5281/zenodo.18677159. The raw data generated in this study can be found by the NCBI GEO accession number GSE319514. The source data for publicly available scRNAseq datasets used in this study can be accessed by 1) NCBI GEO: 9 dpf wildtype Zebrafish Müller glia GSE218107 (GSM6734714), Adult Zebrafish GSE226373 (GSM7074106/ GSM7074109), 2) Single Cell Portal – Broad Cell Atlas: [E18 chick Müller glia, A Cell Atlas of the Chick Retina Based on Single Cell Transcriptomics](https://singlecell.broadinstitute.org/single_cell/study/SCP1159/a-cell-atlas-of-the-chick-retina-based-on-single-cell-transcriptomics?label=AC-1&cluster=Chick%20Muller%20glia%20E18&spatialGroups=--&annotation=Cluster--group--study&subsample=all), and 3) CELLXGENE: [Single cell atlas of the human retina](https://cellxgene.cziscience.com/collections/4c6eaf5c-6d57-4c76-b1e9-60df8c655f1e), [Human fetal retina atlas](https://cellxgene.cziscience.com/collections/c11009b8-b113-4a99-9890-78b2f9df9d79), [Unified comprehensive single-cell atlas of the mouse retina](https://cellxgene.cziscience.com/collections/a0c84e3f-a5ca-4481-b3a5-ccfda0a81ecc).

[dataset] Storey, S. S., Hehr, C. L., Standing, S., McFarlane, S.; 2026; Single cell atlas of the 5 dpf Zebrafish retina; NCBI GEO; GSE319514.

[dataset] Storey, S. S., Hehr, C. L., Standing, S., McFarlane, S.; 2026; Müller glia subtypes define neuro-glial associations and spatial morphogen axes in the zebrafish retina; Zenodo; https://doi.org/10.5281/zenodo.18677159

