## Supplementary Figure 1 for "Müller glia subtypes define neuro-glial associations and spatial morphogen axes in the zebrafish retina"

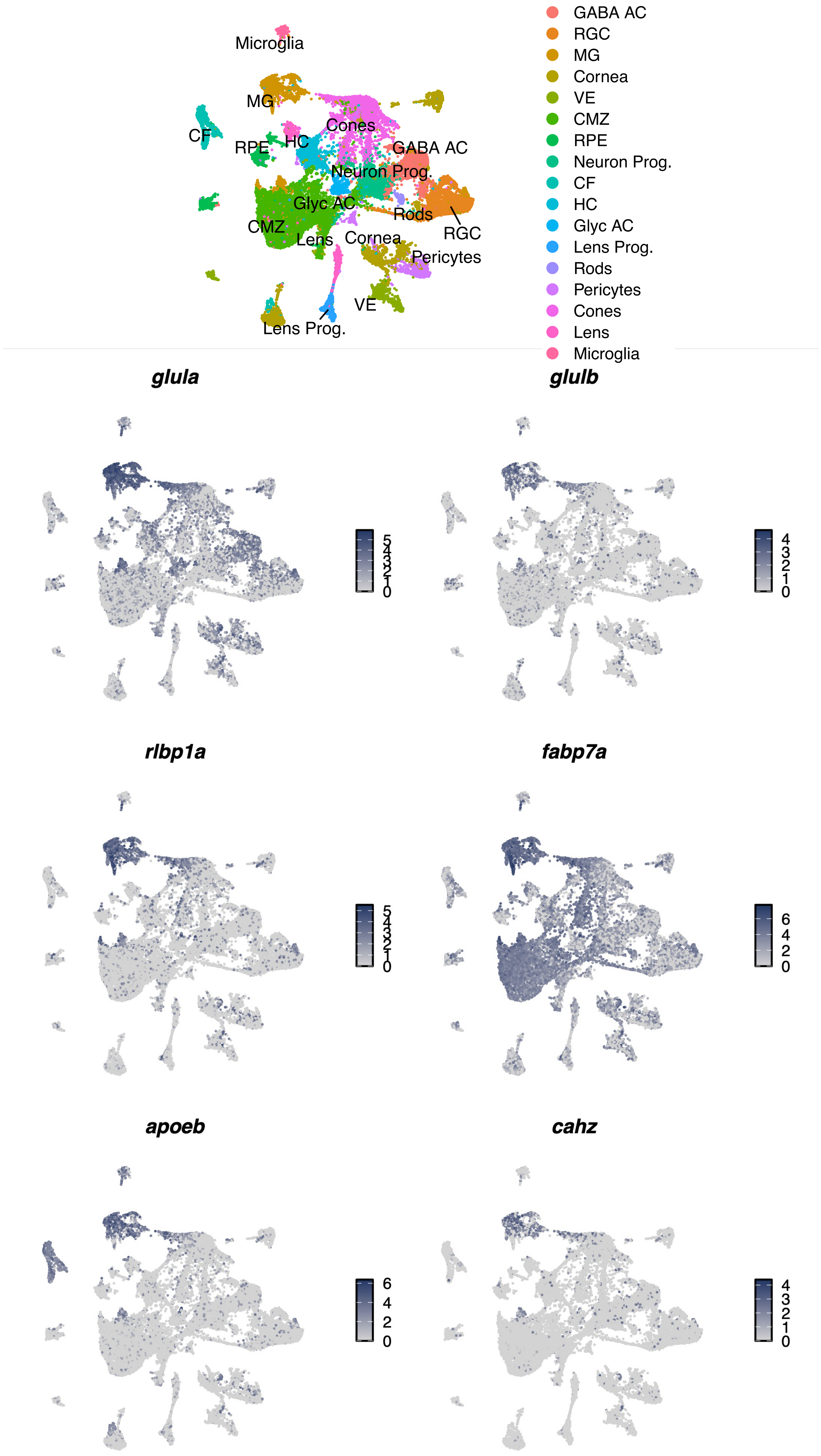


**Supplementary Figure S1. UMAP expression plots for key Müller glia markers in the whole eye of the 5 dpf zebrafish retina.**
