## Supplementary Figure 2 for "Müller glia subtypes define neuro-glial associations and spatial morphogen axes in the zebrafish retina"

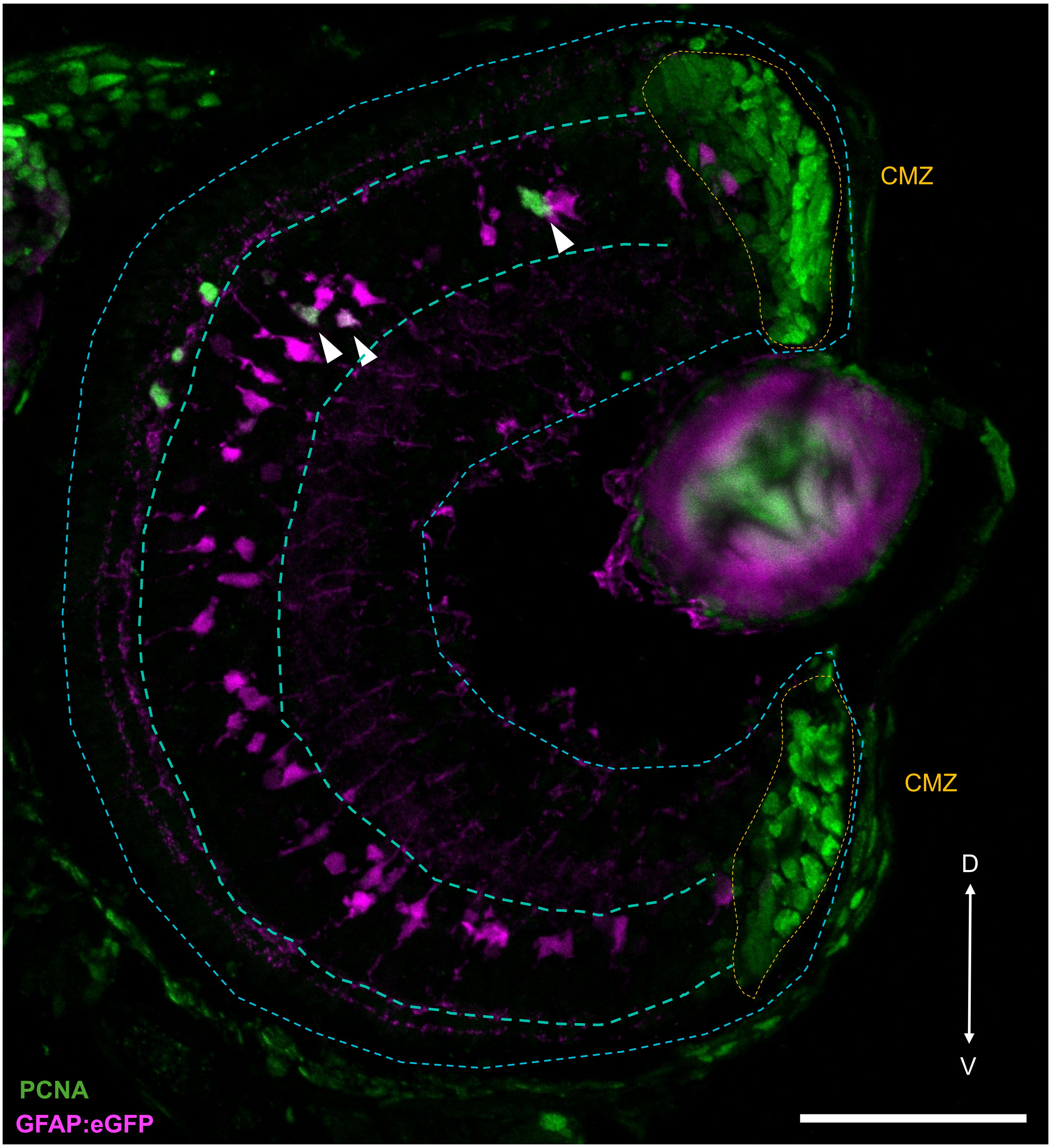


**Supplementary Figure S2. PCNA-immunoreactive Müller glia at the central retina**. Fluorescence immunohistochemistry (IHC) against proliferating cell nuclear antigen (PCNA) in a transverse section of the neural retina of a 5 dpf *Tg(gfap:egfp)* zebrafish. Inner nuclear layer (INL) is shown by blue dashed line. Arrowheads show PCNA+/eGFP+ double positive Müller glia. Axes: D = dorsal, V = ventral. Scale bar: 50 µm.
