## Supplementary Figure 3 for "Müller glia subtypes define neuro-glial associations and spatial morphogen axes in the zebrafish retina"

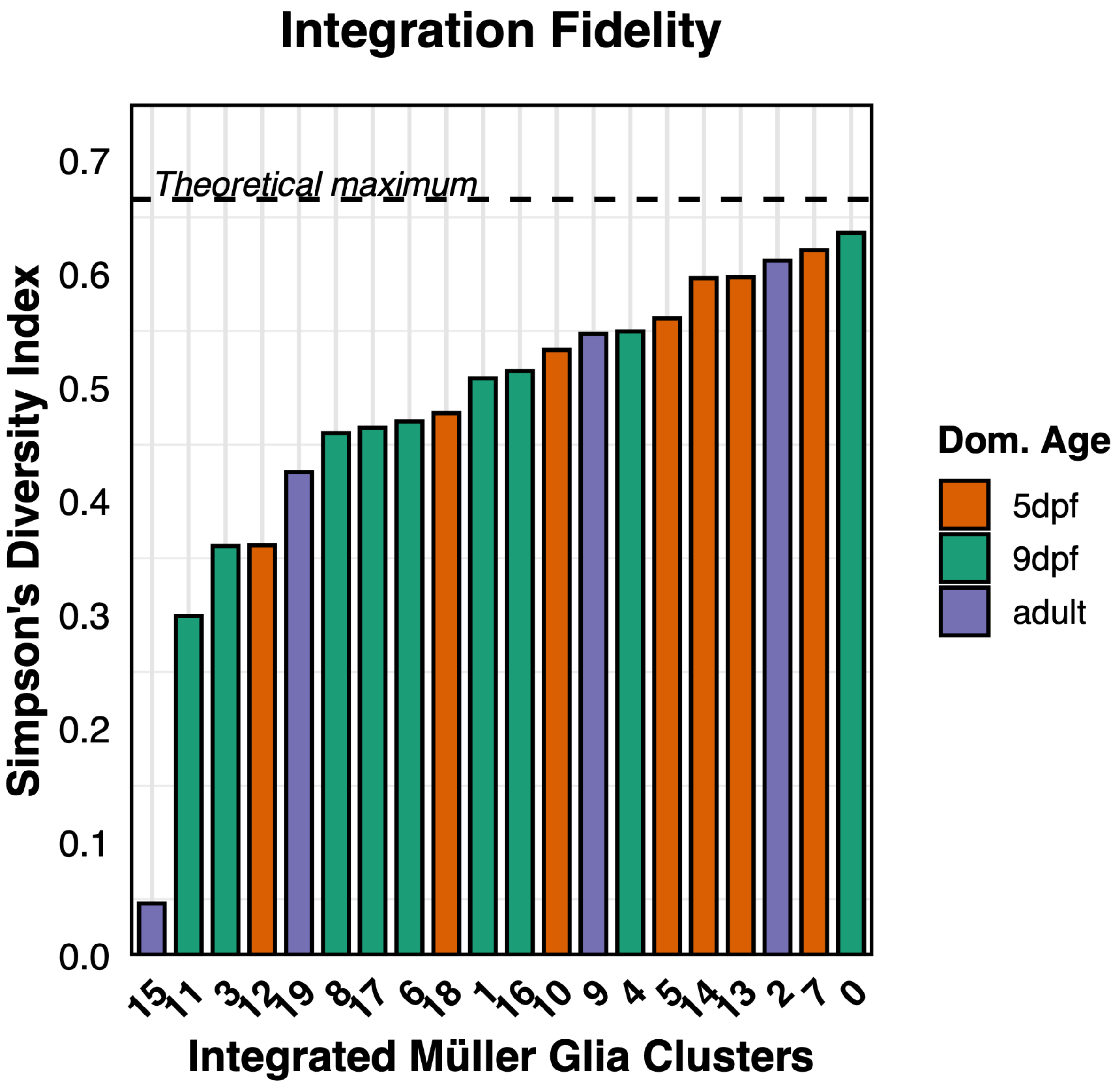


**S3. Barchart of Simpson's diversity index to indicated fidelity of 5dpf, 9dpf and adult zebrafish dataset integration.** Bars colour represents the dominant age for that Cluster.
